# Rooting the animal tree of life

**DOI:** 10.1101/2020.10.27.357798

**Authors:** Yuanning Li, Xing-Xing Shen, Benjamin Evans, Casey W. Dunn, Antonis Rokas

## Abstract

There has been considerable debate about the placement of the root in the animal tree of life, which has emerged as one of the most challenging problems in animal phylogenetics. This debate has major implications for our understanding of the earliest events in animal evolution, including the origin of the nervous system. Some phylogenetic analyses support a root that places the first split in the phylogeny of living animals between sponges and all other animals (the Porifera-sister hypothesis), and others find support for a split between comb jellies and all other animals (Ctenophora-sister). These analyses differ in many respects, including in the genes considered, species considered, molecular evolution models, and software. Here we systematically explore the rooting of the animal tree of life under consistent conditions by synthesizing data and results from 15 previous phylogenomic studies and performing a comprehensive set of new standardized analyses. It has previously been suggested that site-heterogeneous models favor Porifera-sister, but we find that this is not the case. Rather, Porifera-sister is only obtained under a narrow set of conditions when the number of site-heterogeneous categories is unconstrained and range into the hundreds. Site-heterogenous models with a fixed number of dozens of categories support Ctenophora-sister, and cross-validation indicates that such models fit the data just as well as the unconstrained models. Our analyses shed light on an important source of variation between phylogenomic studies of the animal root. The datasets and analyses consolidated here will also be a useful test-platform for the development of phylogenomic methods for this and other difficult problems.

## Main

Historically, there was little debate about the root of the animal tree of life. Porifera-sister (Fig. 1E), the hypothesis that the animal root marks the divergence of Porifera (sponges) from all other animals (Fig. 1B), was widely accepted though rarely tested. By contrast, there has long been uncertainty about the placement of Ctenophora (comb jellies) (Fig. 1D) in the animal tree of life^1^. The first phylogenomic study to include ctenophores^2^ suggested a new hypothesis, now referred to as Ctenophora-sister, that ctenophores rather than sponges are our most distant living animal relative (Fig. 1A). Since then, many more phylogenomic studies have been published (Fig. 2), with some analyses finding support for Ctenophora-sister, some for Porifera-sister, and some neither^3^ (Extended Data Table 1, Supplementary Information section S1). The extensive technical variation across these studies has been important to advancing our understanding of the sensitivity of these analyses, demonstrating for example that outgroup and model selection can have a large impact on these results (Fig. 3A). But the extensive technical variation has also made it difficult to synthesize these results to understand the underlying causes of this sensitivity. Several factors make resolution of the root of the animal tree a particularly challenging problem. For one, the nodes in question are the deepest in the animal tree of life. Another factor that has been invoked is branch lengths *(e.g.* Extended Data Figs. 1,2), which are impacted by both divergence times and shifts in rate of evolution. Some sponges have a longer root to tip length, indicating an accelerated rate of evolution in those lineages. The stem branch of Ctenophora is longer than Porifera stem which, together with a more typical root-to-tip distance, is consistent with a more recent radiation of extant ctenophores^4^ than extant poriferans. The longer ctenophore stem has led some to suggest that Ctenophora-sister could be an artifact of long-branch attraction to outgroups^5^.

**Fig. 1.**
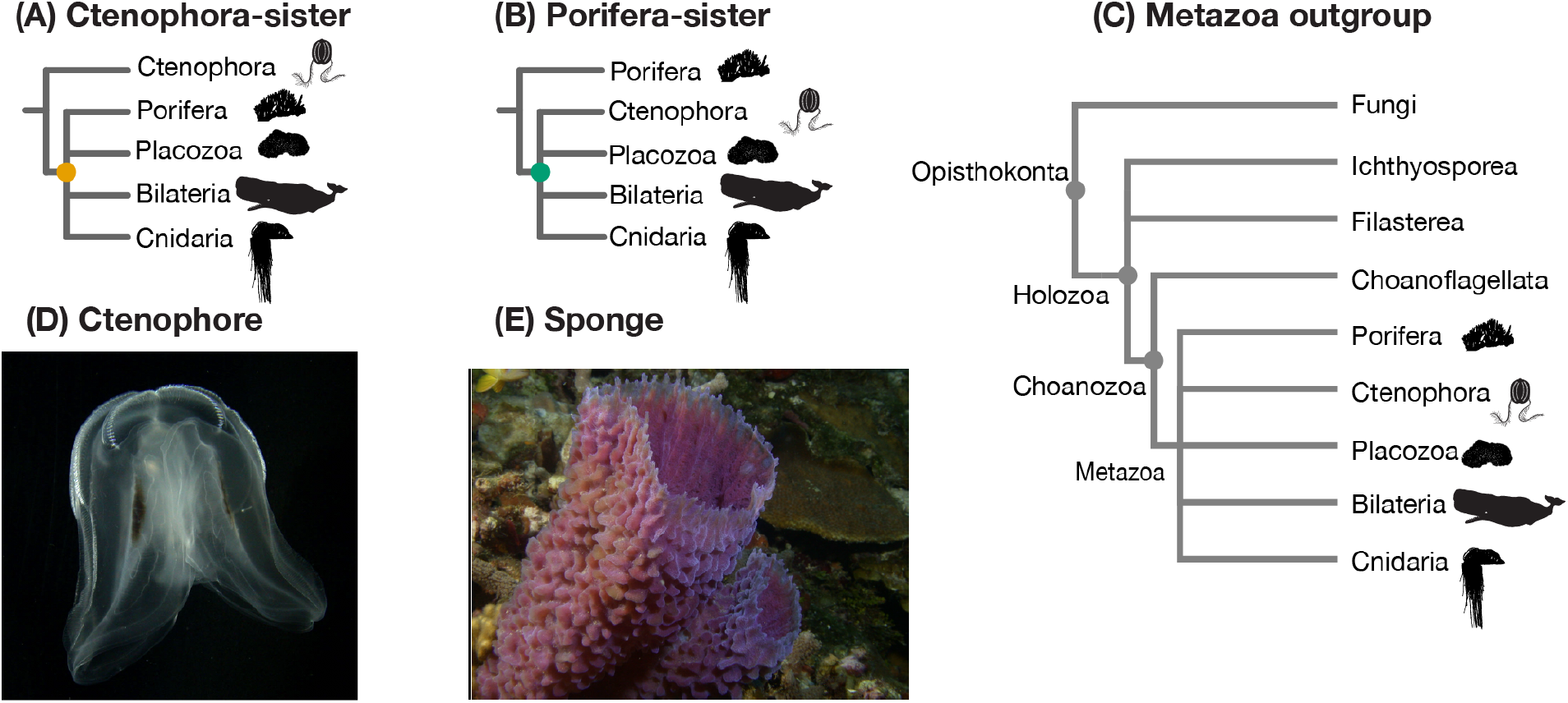
Two alternative phylogenetic hypotheses for the root of the animal tree. (A) The Ctenophora-sister hypothesis posits that there is a clade (designated by the orange node) that includes all animals except Ctenophora, and that Ctenophora is sister to this clade. (B) The Porifera-sister hypothesis posits that there is a clade (designated by the green node) that includes all animals except Porifera, and that Porifera is sister to this clade. Testing these hypotheses requires evaluating the support for each of these alternative nodes. (C) The animals and their outgroup choice, showing the three progressively more inclusive clades Choanozoa, Holozoa, and Opisthokonta. (D) A ctenophore, commonly known as a comb jelliy. (E) A poriferan, commonly known as a sponge.

To advance this debate it is critical to examine variation in methods and results in a standardized and systematic framework. Here, we synthesize data and results from 15 previous phylogenomic studies that tested the Ctenophora-sister and Porifera-sister hypotheses (Fig. 2, Extended Data Table 1). This set includes all phylogenomic studies of amino acid sequence data published before 2019 for which we could obtain data matrices with gene partition annotations. Among these 15 studies, three studies^5,6,7^ are based entirely on previously published data, and gene-partition data is not available from one study^8^.

**Fig. 2.**
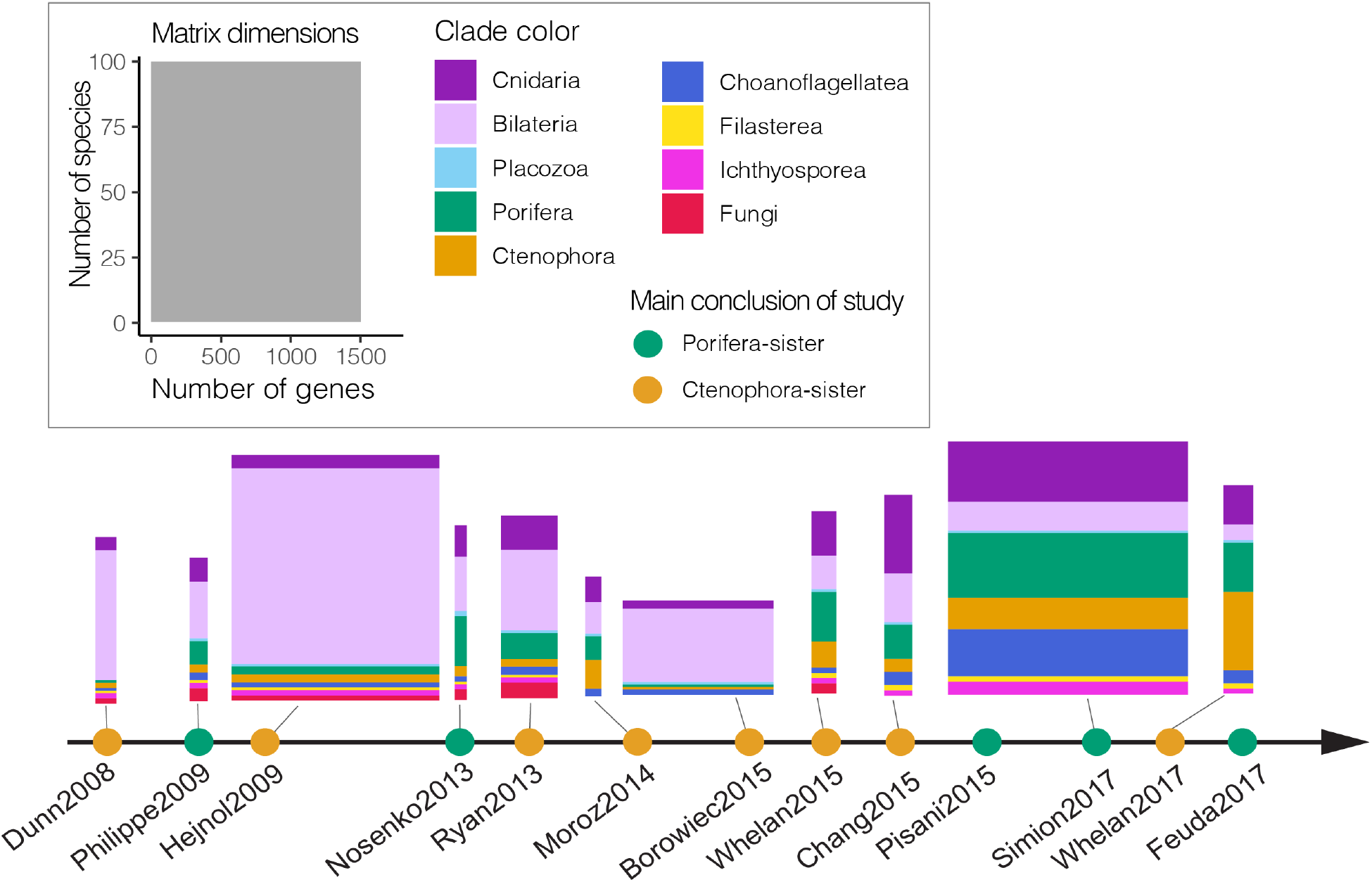
An overview of previous phylogenomic studies on animals. The horizontal axis is a time-series showing the main conclusions of previous phylogenomic studies that indicated their analyses supported Ctenophora-sister (orange nodes) or Porifera-sister (green nodes) based on their main conclusion from each study. Each of the primary matrices considered here is shown above the axis, color coded by taxon sampling.

### Variation in models and sampling across published analyses

The models of sequence evolution in the studies considered here differ according to two primary components: the exchangeability matrix *R* and amino acid equilibrium frequencies Π. The exchangeability matrix *R* describes the relative rates at which one amino acid changes to another. The studies considered here use exchangeabilities that are the same between all amino acids (Poisson, also referred to as F81), or different. If different, the exchangeabilities can either be fixed based on previously empirically estimated rates (WAG or LG), or independently estimated from the data (GTR). The analyses considered here have site-homogeneous exchangeability models (site-homogeneous model), which means that the same matrix is used for all sites.

The equilibrium frequencies describe the expected frequency of each amino acid, which captures the fact that some amino acids are much more common than others. The published analyses differ in whether they take a homogeneous approach and jointly estimate the same frequency across all sites in a partition, or add parameters that allow heterogeneous equilibrium frequencies that differ across sites. Heterogeneous approaches include CAT^9^, which is implemented in the software PhyloBayes and has been widely applied to questions of deep animal phylogeny. The models that are applied in practice are heavily influenced by computational costs, model availability in software, and convention. While studies often discuss CAT and WAG models as if they are mutually exclusive, we note that these particular terms apply to nonexclusive model components – CAT refers to heterogeneous equilibrium frequencies across sites and WAG to a particular exchangeability matrix. In this literature, CAT is generally shorthand for Poisson+CAT and WAG is shorthand for WAG+homogeneous equilibrium frequency estimation. To avoid confusion on this point, here we always specify the exchangeability matrix first *(e.g.* GTR), followed by modifiers that describe the accommodation of heterogeneity in equilibrium frequencies *(e.g.* CAT). If site homogeneous equilibrium frequencies are used, we refer to the exchangeability matrix alone. Gamma-rate heterogeneity, a scalar that accommodates the total rate of change across sites, is used in almost every analysis conducted here and we generally omit its designation. Some analyses partition the data by genes, and use different models for each gene (Supplementary Information section S2).

High-throughput sequencing allows investigators to readily assemble matrices with hundreds or thousands of protein-coding genes from a broad diversity of animal species (Extended Data Table 1). Studies of animal phylogeny have used a wide variety of different approaches to identifying and selecting genes and taxa for their matrices. As a result, the particular genes selected for analysis vary widely (Fig. 2, Extended Data Fig. 3). Gene sampling varies in several ways, including in the fractions of single-copy orthologs *(e.g.* BUSCO genes) and ribosomal protein genes in the matrix (Extended Data Fig. 4).

Horizontal size is proportional to the number of genes sampled, vertical size to the number of species sampled. Only 11 matrices were shown here since three studies^5,6,7^ are based entirely on previously published data and gene-partition data is not available from one study^8^.

Ingroup taxon sampling also varies widely between studies (Fig. 2, Extended Data Fig. 3). Sampling of ingroup taxa (animals) in early studies was biased toward Bilateria. Sampling of non-bilaterian animals, including sponges and ctenophores, has improved over time (Extended Data Fig. 3). Within each clade, there is often considerable variation in taxon sampling and therefore often little species overlap across studies (Extended Data Fig. 3). This variation is in part because newer sequencing technologies in more recent studies are usually not applied to the exact same species that were included in earlier studies.

Sampling of outgroup taxa (non-animals, in this case) is critical to phylogenetic rooting questions, since the node where the outgroup subtree attaches to the ingroup subtree is the root of the ingroup. There has therefore been extensive focus on improving outgroup sampling when testing phylogenetic hypotheses about rooting^10^. Most studies addressing the animal root have removed more distantly related outgroup taxa in some analyses to explore the effect of outgroup selection to ingroup topology^11,5^. Three progressively more inclusive clades have often been investigated: Choanozoa (animals plus most closely related Choanoflagellatea), Holozoa (Choanozoa plus more distantly related Holozoa), and Opisthokonta (Holozoa + Fungi).

### Variation in results across published analyses

We parsed 136 previous phylogenetic analyses from 15 studies (Fig. 3A and Supplementary Table 1). The conclusions of five studies strongly favor Porifera-sister and ten favor Ctenophora-sister (Extended Data Table 1). Three studies are based entirely on previously published data, and the remainder add data for one or more species.s

**Fig. 3.**
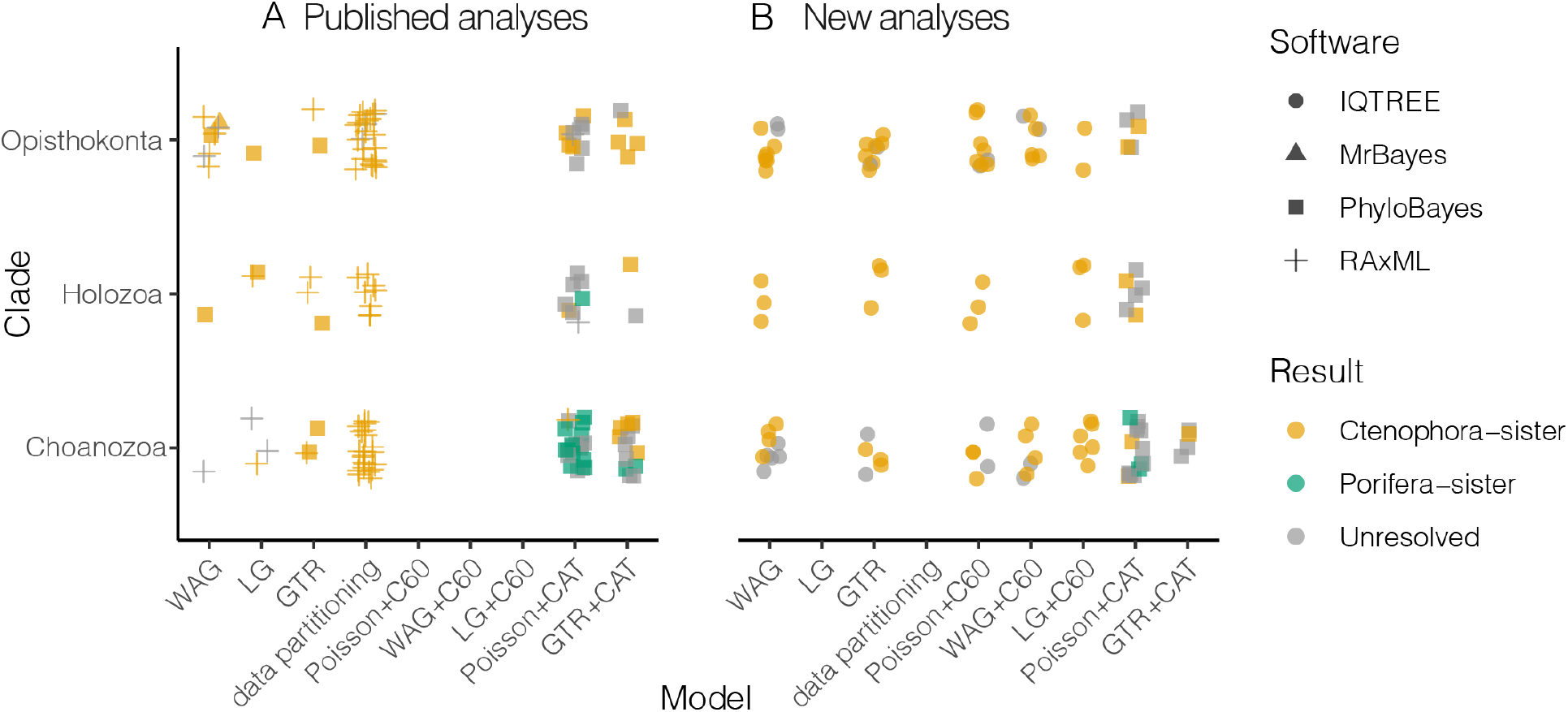
Summary of phylogenomic results from previous studies and reanalysis conducted in this study. Each point represents a phylogenetic analysis with a support of bootstrap values less than 90% or a posterior probability less than 95% are considered as unresolved. The clades are organized with increased outgroup sampling higher on the vertical axis, and the models are organized in general by increasing complexity in terms of numbers of parameters to the right on the horizontal axis. (A) Analyses transcribed from the literature (related to Supplementary Table 1). (B) New phylogenomic analyses conducted in this study (related to Supplementary Table 2). Over 1,011,765 CPU hours (~115 CPU years) were used for this study.

Our summary of previous phylogenetic analyses (Fig. 3A) shows that Ctenophora-sister is supported in analyses that span the full range of outgroup sampling and models used to date, including some analyses with restricted outgroup sampling and models that accommodate site-heterogeneous equilibrium frequencies with CAT. This is consistent with previous assessments of the problem^6^, but is drawn from a much more extensive and systematic examination. The only analyses that support Porifera-sister have reduced outgroup sampling (Choanozoa, Holozoa) and site-heterogeneous models with CAT. Model adequacy assessments generally favor GTR+CAT over Poisson+CAT or site-homogeneous models^5,7^, but because GTR+CAT is so parameter rich, many analyses that use a model with GTR+CAT do not converge. The fact that Porifera-sister is recovered only for particular models with particular outgroup sampling indicates that model and outgroup interact, and that this interaction is fundamental to understanding the range of results obtained across analyses.

### New standardized analyses of published matrices

One of the challenges of interpreting support for the placement of the animal root across published studies is that different programs, software versions, and settings have been used across analyses. This extensive variation makes it difficult to identify the primary factors that lead to different results. Here we first reanalyze the primary matrices from each study under the same conditions with IQ-TREE^12^ with multiple evolutionary models. We selected this tool because it has greater model flexibility than most other phylogenetic tools^13^ (Fig. 3B; Supplementary Table 3). Importantly, it has C models (C10-C60 equilibrium frequencies)^14^ that, like CAT, allow models to accommodate heterogeneity in equilibrium amino acid frequencies across sites.

For each of the published studies, we selected the matrix that was the primary focus of the manuscript, or has been reanalyzed extensively in other studies, for further analysis. For each of these matrices, we progressively trimmed taxon sampling to create Opisthokonta, Holozoa, and Choanozoa versions, where permitted by original outgroup sampling. This produced 36 data matrices from 11 studies that presented new sequence data and for which partition data were available.

For all but the three largest matrices, we tested the relative fit of a variety of models, both with and without C10-C60 accommodation of site heterogeneity in equilibrium frequencies, using ModelFinder^15^ in IQ-TREE. In all cases, models with C60 fit these matrices better than the site-homogeneous models. This is consistent with the importance of accommodating site heterogeneity noted by previous investigators^16,17,18,5,7,9^. We then inferred support under the best-fit model (Supplementary Table 3), except for the three largest matrices where we used LG+C60. We then analyzed each matrix under a panel of standard site-heterogeneous and site-homogeneous models, including WAG, GTR and Poisson+C60 (Supplementary Table 3).

All IQ-TREE analyses, apart from unresolved analyses (for Moroz2014_3d and all Nosenko2013 matrices), supported Ctenophora-sister (Fig. 3B). No IQ-TREE analyses supported Porifera-sister, including those that restrict outgroup sampling to Choanoflagellatea and use models with site-heterogeneous equilibrium frequencies (Fig. 3B lower right; Supplementary Table 2), the conditions under which published PhyloBayes CAT analyses recover strong support for Porifera-sister (Fig. 3A). To further verify this difference in a controlled manner we reran PhyloBayes analyses with CAT, using both Poisson and GTR substitution matrices, for some matrices that had led to support for Porifera-sister in published analyses. Consistent with published results, some of these supported Porifera-sister.

Our new analyses show that, with restricted outgroup sampling, analyses of the same matrices with two different means of accommodating site heterogeneity in equilibrium frequencies (C60 in IQ-TREE and CAT in PhyloBayes) yield different results. This indicates that the traditional framing of the problem, that accommodating site heterogeneity leads to support for Porifera-sister, is not correct. Instead, there is something about the PhyloBayes CAT analyses specifically that leads to support for Porifera-sister.

### Category number explains differences between site-heterogeneous analyses

There are several factors, including variations in models (C60 vs CAT), software (Phylobayes vs IQ-TREE), and implementation details (*e.g.* number of categories used to accommodate site heterogeneity) that could explain the new variation in results noted here among site-heterogeneous models. In published analyses of the animal root, these factors were confounded since all previous heterogeneous analyses used the CAT model in PhyloBayes. Here we seek to deconfound these factors to gain a finer-grained perspective on why results differ between analyses of the same matrices.

A primary difference between the C (like C60) and CAT site-heterogeneous models is the number of equilibrium frequency categories. The standard CAT model employs a Dirichlet process prior to inferring the number of equilibrium frequency categories, so the number of categories is variable^9^. IQ-TREE implements C models^14^ with a fixed number of categories, from 10 (C10) to 60 (C60). Differences in analysis results could therefore be due to differences in the number of categories. The number of categories inferred by CAT in PhyloBayes can be very high (Supplementary Table 4), with a mean here of 623.5 categories for Poisson+CAT analyses in all matrices and 1026 categories for GTR+CAT analyses in several representative matrices. This requires a very large number of additional estiamted parameters.

We examined the specific impact of this large difference in category number on animal root position. It is currently not possible to use more than 60 categories for C models in IQ-TREE, but the number of categories can be set *a priori* in Phylobayes using nCAT. We therefore varied the number of categories in Phylobayes analyses (Fig. 4; Supplementary Table 5). We found that Poisson+nCAT60 analyses in PhyloBayes, like Poisson+C60 and WAG+C60 IQ-TREE analyses, provide strong support of Ctenophora-sister. This indicates that the difference in results between unconstrained CAT analyses in PhyloBayes and C60 analyses in IQ-TREE is not due to differences in the software or other implementation factors, but due to the large difference, in excess of ten-fold, in the number of site categories. When we increased the number of categories in PhyloBayes nCAT analyses, we observe the transition from support for Ctenophora-sister to an unresolved root to Porifera-sister (Fig. 4). For example, for the Whelan2017_strict matrix this transition occurs between 60 – 120 categories when using Poisson+nCAT model. Due to computational limitations of GTR models, we only ran GTR+CAT and GTR+nCAT60 models on representative matrices with Choanozoa sampling. We found that for Whelan2017 matrices, support shifted from Porifera-sister with Poisson+CAT model to Ctenophora-sister using GTR+CAT model. Moreover, we also found all results strongly supported Ctenophora-sister with GTR+nCAT60 models.

**Fig. 4.**
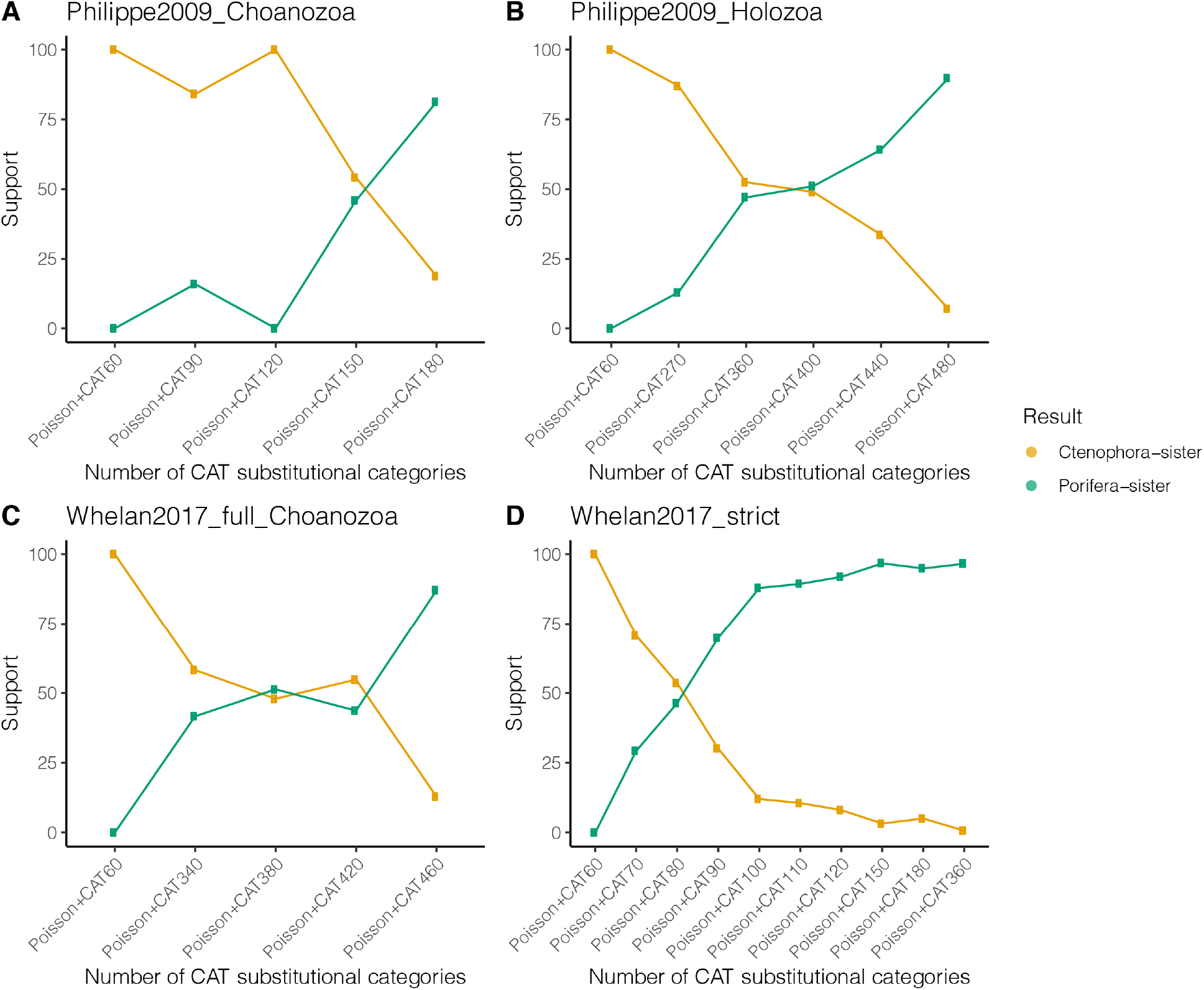
Sensitivity analyses with representative data matrices were analyzed by different number of equilibrium frequency categories (nCAT) in PhyloBayes. Statistical support values (posterior probabilities) were obtained from three data matrices using the site-heterogeneous Poisson+CAT model with different categories. (A) Phlippe2009_Choanozoa; (B) Philippe2009_Holozoa; (C) Whelan2017_full_Choanozoa; (D) Whelan2017_strict. Statistical support for Ctenophora-sister and Porifera-sister is indicated in orange and green, respectively. Support values from the sensitivity analyses are shown in Supplementary Table 5.

These results further clarify when analyses support Porifera-sister (*e.g.* Whelan2017: Extended Data Fig. 1; Philippe2009: Extended Data Fig. 2): when outgroup sampling is restricted (Choanozoa), when a Poisson (rather than a GTR) exchange matrix is used, and when a very large number of site categories is used (unconstrained CAT, giving hundreds of equilibrium frequency categories). Analyses under other conditions either support Ctenophora-sister or are unresolved. This is consistent across published analyses and our new panels of analyses. The question, then, isn’t why similar analyses give different results, but how we should interpret variation in results when we run analyses that differ in these specific respects. If we fix the first two features to conditions that are necessary for Porifera-sister support (Choanozoa taxon sampling and a Poisson exchange matrix), there are several insights that we can glean from examining how the number of equilibrium frequency categories impacts results that sheds light on interpretation of those results.

The first questions regard model fit. ModelFinder selects site heterogeneous C60 models according to BIC (Supplementary Table 3), but IQ-TREE often gives a warning under C60 that the model may overfit, with too many categories, because some mixture weights are close to 0 (Supplementary Table 6). In Phylob-Bayes, Cross-validation is a reliable and suggested approach to model evaluation^19^. We elvaluated Pois-son+CAT and Poisson+nCAT60 models with cross-validation in PhyloBayes for the Whelan2017_strict and Philippe2009_Choanozoa matrices. For both matrices, we found nearly identical distributions of cross-validation scores for Poisson+nCAT60 and Poisson+CAT models (Extended Data Fig. 5). A t-test analysis shows that there is no significant difference between them for either matrix (Whelan2017_strict: *p — value* = 0.86; Philippe2009_Choanozoa: *p — value* = 0.99). Cross-validation therefore does not indicate that the unconstrained CAT models are a better fit than the models with 60 categories. This suggests there is not a reason to favor the narrow analysis conditions (Choanozoa taxon sampling, Poisson exchange matrix, and unconstrained CAT models that use hundreds of categories) under which Porifera-sister are obtained.

In a site-heterogeneous model, each site is allocated to one of the categories in each sample. Different categories can have very different numbers of sites. We examine category allocations in the last chain samples of analyses of the Philippe2009_Choanozoa and Whelan2017_strict matrices (Extended Data Fig. 6A-B), which have 510 and 471 categories, respectively. The fraction of sites allocated to the 50% of the least frequent categories is 2.97% in the analysis of the Philippe2009_Choanozoa matrix and 3.46% in the analysis of the Whelan2017_strict matrix. This long tail of categories that apply to a very small fraction of sites is in stark contrast to nCAT60 analyses, constrained to 60 categories, which have no such long tail of rare categories (Extended Data Fig. 6A-B). The fact that so many categories apply to such a small fraction of data may help explain why increasing the number of categories more than ten-fold has so little impact on the predictive power of these far more complex models.

### Phylogenetic signal

To further explore the phylogenetic signal of different models we discussed above, we quantified the phylogenetic signal for Porifera-sister and Ctenophora-sister topologies across three representative data matrices when varying outgroup sampling and model (Extended Data Fig. 7). By calculating differences in loglikelihood scores for these topologies for every gene (△|lnL|) in each matrix when using site-homogeneous models in IQ-TREE, we found that the Ctenophora-sister had the higher proportions of supporting genes in every analysis. Moreover, outgroup choice has little impact on the distribution of the support for phylogenetic signals in analyses with site-homogeneous models. This finding is largely consistent with the previously observed distribution of support for Ctenophora-sister in other data matrices^20^.

Although a higher proportion of genes support Ctenophora-sister with site-heterogeneous C60 models, the phylogenetic signal decreases in many genes using C60 models compared to site-homogeneous models. In an extreme case, in matrices from Ryan2013_est nearly 30% of genes changed from strong ∆|lnL|>2) to weak Ctenophora-sister signal (∆|lnL|<2) (Extended Data Fig. 7; Supplementary Table 7). In contrast to the C60 models, there is a major increase in phylogenetic signal in Poisson+CAT models in PhyloBayes towards Porifera-sister, and outgroup choice has a major effect of the distribution of phylogenetic signal (Extended Data Fig. 7). For example, in Whelan2017_full matrix, we found that the number of genes that support Ctenophora-sister in analyses with CAT decreases from 57.5% in matrices with distant outgroups (Holozoa) to 35.4% when outgroups are restricted (Choanozoa, Supplementary Table 7).

### Other approaches to accommodating site heterogeneity

Other approaches have been taken to addressing base compositional heterogeneity across taxa. For example, Feuda *et al.*^21^ recoded the full set of twenty amino acids into six groups of amino acids. These groups tend to have more frequent evolutionary changes within them than between them^22^. Recoding could, like CAT and C models, address variation across sites, but it could also accommodate variation across lineages, and it was suggested that this approach favors Porifera-sister^21^. However, our own analyses (Supplementary Information section S3, Supplementary Figs. 8-9) comparing the performance of random and empirical recoding indicate that the impact of data recoding is largely due to discarding information, not accommodating variation in amino acid composition. Consistent with a recent simulation study on data recoding^23^, these findings indicate that recoding can be a problematic method for addressing heterogeneity.

## Conclusions

Resolving the placement of the root in the animal tree of life has proved very challenging^24^. By synthesizing past phylogenomic studies and performing new analyses, we find that support of Porifera-sister is only recovered by site-heterogeneous CAT models with restricted outgroup sampling, and then only in some such analyses. Through controlled analyses we are able to identify the specific aspect of the models that is involved in this variation – the number of categories used to accommodate site heterogeneity in equilibrium frequency (Fig. 4). The 10-fold difference in category number seen in the more complex CAT models that support Porifera-sister does not improve model fit according to cross-validation, though. This suggests that we shouldn’t privilege these narrow analysis conditions that recover Porifera-sister over the much broader range of conditions that recover Ctenophora-sister.

Pin-pointing category number as an issue with large effect on analyses of the animal root will help guide future analyses that address this question. We hope that the work we have conducted here to consolidate many datasets and analyses in standard formats will make it easier for other investigators to engage in this particularly interesting and difficult phylogenetic problem, and that this problem can be a test-bed to develop methods and tools that will help with other difficult phylogenetic problems as well. Advances on the question of the animal root will come from progress on other fronts as well. For example, there are many organisms that are highly relevant to this problem, in particular outgroup, ctenophore, and sponge taxa, for which no genome or transcriptome data are available^3^. More broadly, there are very few chromosome-level genome assemblies for animals outside of Bilateria. Future analyses focused on complete genomes rather than transcriptomes and partial genomes will have multiple advantages. Data matrices derived from these more complete sources will have a lower fraction of missing sequences. Complete gene sampling within each species will also greatly improve analyses of gene duplciation and loss, a crititical step in building phylogenomic matrices such as those presented here. Analyses of this new generation of matrices derived from complete genomes will be well served my understandind the sources of analysis variation in the generation of matrices that came before them.

## Methods

### Data and code availability

The main data and results associated with the main text and supplementary materials are available in the GitHub data repository at https://github.com/dunnlab/animal_tree_root. All tree files, intermediate results and scripts/commands associated with this study are available in the Figshare data repository at https://doi.org/10.6084/m9.figshare.13085081.v1.

### Data selection and wrangling

We retrieved matrices from each publication (Extended Data Table 1), storing the raw data in this manuscript’s version control repository. We manually made some formatting changes to make the batch processing of the matrices work well, *e.g.* standardizing the format of Nexus CHARSET blocks. All changes made are tracked with git.

### Matrix comparison and annotation

#### Taxon name reconciliation

We programmatically queried the NCBI Taxonomy database to standardize names of samples in each matrix. We also used a table where manual entries were needed (manual_taxonomy_map.tsv), *e.g.* authors of the original matrix indicate species name in original manuscript. For a table summarizing all samples and their new or lengthened names, see taxon_table.tsv.

#### Sequence comparisons

Using the original partition files for each matrix, we separated each sequence for each taxon from each partition. Because many of the matrices had been processed by the original authors to remove columns that are poorly sampled or highly variable, these matrix-derived sequences can have deletions relative to the actual gene sequences.

We used DIAMOND v0.9.26^25^ to compare each sequence to all others using default diamond Blastp parameters. We further filtered DIAMOND results such that we retained hits for 90% of partitions (pident > 50.0, eValue < 1e-5, no self vs self). We ran BUSCO with default parameters for all sequences against the provided Metazoa gene set. We also ran a BLAST+ v2.8.1^26^ blastp search against the SwissProt^27^ database, filtering results such that we retain at least one hit for ~97% of partitions (pident > 50.0, eValue < 1e-15).

#### Partition network

We used the sequence similarity comparisons described above to compare partitions.

We constructed a network with Python and NetworkX v2.2^28^ where each node is a partition and each edge represents a DIAMOND sequence-to-sequence match between sequences in the partitions. We extracted each connected component from this network. We further split these components if the most connected node (i.e. most edges) had two times more the standard deviation from the mean number of edges in the component it is a member of and if removing that node splits the component into two or more components. We then decorated every node in the partition network with the most often found SwissProt BLAST+ result and BUSCO results to see which components contain which classes and families of genes. See partition_network_summary in Rdata for a summary tally of each part of the comparison.

### Phylogenetic analyses

#### Phylogenetic analyses in IQ-TREE

To investigate the phylogenetic hypotheses and distribution of phylogenetic signal in studies aiming to find the root position of animal phylogeny, we considered 16 data matrices from all phylogenomic studies that were constructed from EST, transcriptomic, or genomic data (Extended Data Table 1). Because different choices of substitution models could largely influence phylogenetic inference of the placement of the root position of animal phylogeny *(e.g.* site-heterogeneous vs. site-homogeneous models), we first investigated model-fit from each matrix using ModelFinder in IQ-TREE v1.6.7, including site-heterogenous C10 to C60 profile mixture models (C60 models) as variants of the CAT models in ML framework (C10-C60 model were included for model comparison via -madd option). We included models that are commonly used in previous analyses, including site-homogeneous Poisson, WAG, LG, GTR models plus C10-C60 models in the model testing. For computational efficiency, the GTR+C60 models were not included in model testing since it requires to estimate over 10,000 parameters. For large matrices like those from Hejnol2009, Borrowiec2015, and Simion2017, model testing is also not computational feasible so only LG+C60 models were used since LG/WAG+C60 models were suggested as the best-fit model in all other matrices.

We then reanalyzed each matrix under a panel of evolutionary models, including WAG, GTR, Poisson+C60 and associated best-fit model identified above. Nodal support was assessed with 1000 ultrafast bootstrap replicates for each analysis. Because of the large size of Hejnol2009 and Simion2017, it was not computationally feasible to analyze the whole matrix using the C60 model or CAT site-heterogeneous models. To circumvent his limitation, we reduced the data size from their full matrices to facilitate computational efficiency for site-heterogeneous models. For Hejnol2009 matrix, we instead used the 330-gene matrix constructed by Hejnol *et al.* 2009, since the main conclusion for their study is based on this subsampled matrix; For Simion2017 matrix, we only included the most complete 25% of genes (genes that were present in less than 79 taxa were removed; 428 genes were kept). It should be noted that the main conclusion of Simion *et al*. was also based on selection of 25% of genes for their jackknife approach.

#### Outgroup taxa sampling with C60 and CAT models

Because different choices of outgroups could also affect phylogenetic inference as suggested in previous analyses, we parsed the full data matrices into three different types of outgroups: Choanozoa, Holozoa and Opisthokonta. These datasets include the same set of genes but differ in the composition of outgroup species. Choanozoa only includes Choanoflagellatea outgroup; Holozoa also includes more distantly related Holozoans; Opistokonta also includes Fungi. For each Choanozoa data matrice, both C60 models in IQ-TREE and Poisson+CAT models in PhyloBayes were conducted. The maximum likelihood analysis was performed using the best-fit substitution model identified as above and nodal support was assessed with 1000 ultrafast bootstrap replicates using IQ-TREE. Moreover, Bayesian inference with the site-heterogeneous Poisson+CAT model was done with PhyloBayes-MPI v1.8. To minimize computational burden, GTR+CAT models were only performed in the representative Choanozoa matrices from Philippe2009, Ryan2013_est and Whelan2017_full.

For several Choanozoa matrices indicated strong support for the hypothesis that sponges are the sister group to the remaining Metazoa using the Poisson+CAT model, Bayesian inference with Poisson+CAT model was also conducted to Holozoa and Opisthokonta data matrices with the same settings as above. For all the analyses with Poisson+CAT models in PhyloBayes, two independent chains were sampled every generation. Tracer plots of MCMC runs were visually inspected in Tracer v1.6 to assess stationarity and appropriate burn-in. Chains were considered to have reached convergence when the maxdiff statistic among chains was below 0.3 (as measured by bpcomp) and effective sample size > 50 for each parameter (as measured by tracecomp). A 50% majority-rule consensus tree was computed with bpcomp, and nodal support was estimated by posterior probability. Most Phylobayes runs converged, although several large matrices have not reached convergence after at least a month’s computational time. For those matrices that were not converged, PhyloBayes analyses were run for at least two weeks. We also summarized the average number of substitutional categories inferred for each PhyloBayes analysis using Tracer.

#### Phylogenetic distribution of support

To investigate the distribution of phylogenetic signal of the animal-root position in data matrices, we considered three major data matrices from three studies that had different topology between ML and BI using CAT model in our reanalysis, including Philippe2008, Ryan2013_est, and Whelan2017_full data matrices. We examined two hypotheses: Ctenophora-sister (T1) and Porifera-sister (T2) to the rest of metazoans, under a panel of evolutionary models with different outgroup schemes (Choanozoa and the full matrix). For IQ-TREE analysis in each data matrix, site-wise likelihood scores were inferred for both hypotheses using IQ-TREE (option -g) with the LG+G4 model. The two different phylogenetic trees passed to IQ-TREE 1.6.12 (via -z) were the tree where Ctenophora-sister and a tree modified to have Porifera placed as the sister to the rest of animals. The numbers of genes and sites supporting each hypothesis were calculated from IQ-TREE output and Perl scripts from a previous study^20^. By calculating gene-wise log-likelihood scores between T1 and T2 for every gene, we considered a gene with an absolute value of log-likelihood difference of two as a gene with strong (|ΔlnL|> 2) or weak (|ΔlnL|< 2) phylogenetic signal as done in a previous study^29^.

For Poisson+CAT and LG in PhyloBayes, we only considered the Philippe2009 and Whelan2017 matrices due to computational efficiency. Since the default option in PhyloBayes does not provide the feature to calculate site-wise log likelihood for every generation, we replaced the line “int sitelogl = 0” with “int sitelogl = 1” in the file named “ReadSample.cpp” and installed PhyloBayes 4.1c so that site-wise log likelihood value can be stored to a file that ends with “.sitelogl” (via readpb -sitelogl). For each condition, we first calculated site-wise log likelihoods for each of two hypotheses (T1 and T2) using pb (via -T) and then stored site-wise log likelihood (a total number of samples for each site is 20) every ten until 300th generations, after discarding the first 100 generations using readpb (via -sitelogl -x 100 10 300). Next, we normalized site-wise log likelihood value across 20 samples for each of two hypotheses (T1 and T2) and combined normalized sitewise log likelihood values of T1 and T2 into a single file that was used to calculate gene-wise log-likelihood scores between T1 and T2 with Perl scripts from a previous study^20^.

#### Sensitivity analyses with different number of substitutional categories

To explore how the number of substitutional categories may affect the phylogenetic inference related to the animal phylogeny, we conducted PhyloBayes analyses with a panel of different substitutional categories in the Whelan2017_strict (ncat=60, 70, 80, 90, 110, 120, 150, 180, 360), Whelan2017_full_Choanozoa (nCAT=60, 340, 380, 420, 460), Philippe2009 (nCAT=60, 90, 120, 150, 180) Philippe2009_Holozoa (nCAT=360, 400, 440, 480) and Ryan2013_est (ncat=60). To compare the results between Poisson+CAT and GTR+CAT and minimize computational burden, GTR+CAT and GTR+CAT60 models were only performed in the representative Choanozoa matrices from Philippe2009, Ryan2013_est and Whelan2017_full. All PhyloBayes analyses were carried out using the same settings as above (see Outgroup taxa sampling with C60 and CAT models section), except when a different number of categories was used.

To compare the allocation of frequency categories across sites in the Philippe2009_Choanozoa and Whe-lan2017_strict matrices for the constrained Poisson+CAT60 model and unconstrained Posson+CAT model, we parsed the information of PhyloBayes chain files by sampling one in every 1000 generations after burnin determined above. The scripts and subsampled chain files are in the `../trees_new/frequency` subdirectory of the git repository.

#### Cross-validation analyses

Bayesian cross-validation implemented in PhyloBayes-MPI was used to compare the fit of Poisson+nCAT60 and Poisson+CAT models coupled with a gamma distribution of site-rate heterogeneity in Whe-lan2017_strict and Philippe2009_Choanozoa data matrices. Ten replicates were run, each replicate consisting of a random subsample of 10,000 sites for training the model and 2,000 sites for computing the cross-validation likelihood score. For each run, 5,000 generations were run and the first 2,000 generations were discarded as burn-in.

#### Performance of data-recoding

All code used for the analyses presented here is available in a git repository at https://github.com/caseywdunn/feuda_2017. The randomized recoding analyses are in the `recoding/alternative` subdirectory of the git repository.

The original SR-6 recoding scheme is “APST CW DEGN FHY ILMV KQR”^22^, where spaces separate amino acids that are placed into the same group. This recoding is one member of a family of recodings, each with a different number of groups, based on clustering of the JTT matrix. The other recoding used by Feuda *et al.,* KGB-6 and D-6, are based on different matrices and methods^21^.

The `alt_recode.py` script was used to generate the randomized recoding schemes and apply the recodings to the data. To create the randomized recoding schemes, the amino acids in the SR-6 recoding were randomly reshuffled. This creates new recodings that have the same sized groups as SR-6. The new recodings were, from `random-00` to `random-03`:

MPKE AY IDGQ TRC WNLF SVH

EIFT WL QVPG NKM SCHA RYD

LCQS GK WPTI VRD YEFN MAH

IWQV TY KDLM ESH APCF GRN

To apply these to the data, each amino acid was replaced with the first amino acid in the group. When applying `random-00`, for example, each instance of `R` and `C` would be replaced by a `T`.

The 20 state matrices are the same across all analyses since they are not recoded. Since all 20 state matrices are the same, variation between 20-state results (as in the left side of each pane of Extended Data Fig. 9) gives insight into the technical variance of the inference process.

Each new matrix was analyzed with PhyloBayes-MPI. Analyses were run for 1000 generations, and a 200 generation burnin applied. The resulting tree files and posterior predictive scores were parsed for display with the code in `manuscript.rmd`.

The statistics presented in Extended Data Fig. 8A were parsed from the Feuda *et al.* manuscript into the file `tidy_table_3.tsv` and rendered for display with the code in `manuscript.rmd`.

## Acknowledgements

We thank members of the Dunn and Rokas laboratories for discussions and comments. We are grateful to Steve Haddock, Kyle David, Stephen Haddock and Nathan Whelan for feedback on an earlier version of this manuscript. We are grateful to Nicole King, Daniel Richter, Paul Lewis, Warren Francis and Joseph Ryan for feedback on the preprint version of this manuscript. We thank the Yale Center for Research Computing for use of the research computing infrastructure, specifically the Farnam HPC cluster. Yuanning Li was partially supported by a scholarship from the China Scholarship Council (CSC) for studying and living abroad. This work was conducted in part using the resources of the Advanced Computing Center for Research and Education (ACCRE) at Vanderbilt University. Research in A.R.’s lab is supported by the National Science Foundation (DEB-1442113), the National Institutes of Health / National Institute of Allergy and Infectious Diseases (1R56AI146096-01A1), the Guggenheim Foundation, and the Burroughs Wellcome Fund.

## Author contributions

C.W.D., Y.L. conceived this study. C.W.D. and A.R. supervised the project. Y.L., B.E., X-X.S., and C.W.D. conducted analyses. Y.L., B.E. and C.W.D. wrote the paper. All authors discussed the results and implications and commented on the manuscript at all stages.

## Extended Tables

**Extended Data Table 1.** Overview of data matrices used in this study.

| Study | Matrix | Clade | No. partitions | No. sites | No. taxa | Inferred sister lineage | Journal | No. citations |
| --- | --- | --- | --- | --- | --- | --- | --- | --- |
| Dunn2008 | Dunn2008 | Opisthokonta | 150 | 21,152 | 64 | Ctenophora-sister | Nature | 1,794 |
| Philippe2009 | Philippe2009 | Opisthokonta | 129 | 30,257 | 55 | Porifera-sister | Current Biology | 649 |
| Hejnol2009 | Hejnol2009 | Opisthokonta | 1,487 | 270,580 | 94 | Ctenophora-sister | Proc. Biol. Sci. | 707 |
|  | Hejnol2009_subsampled | Opisthokonta | 844 | 55,594 | 94 | Ctenophora-sister |  |  |
| Pick2010 |  |  |  |  |  | Porifera-sister | Molecular Biology and Evolution | 326 |
| Nosenko2013 | Nosenko2013_nonribo_9187 | Opisthokonta | 35 | 9,187 | 63 | Ctenophora-sister | Molecular Phylogenetics and Evolution | 221 |
|  | Nosenko2013_ribo_14615 | Opisthokonta | 87 | 14,615 | 71 | Porifera-sister |  |  |
| Ryan2013 | Ryan2013_est | Opisthokonta | 406 | 88,384 | 61 | Ctenophora-sister | Science | 534 |
| Moroz2014 | Moroz2014_3d | Opisthokonta | 114 | 22,772 | 46 | Ctenophora-sister | Nature | 491 |
| Whelan2015 | Whelan2015_D1 | Opisthokonta | 251 | 59,733 | 76 | Ctenophora-sister | PNAS | 220 |
|  | Whelan2015_D10 | Opisthokonta | 210 | 59,733 | 70 | Ctenophora-sister |  |  |
|  | Whelan2015_D20 | Opisthokonta | 178 | 81,008 | 65 | Ctenophora-sister |  |  |
| Borowiec2015 | Borowiec2015_Best108 | Choaimalia | 108 | 41,808 | 36 | Ctenophora-sister | BMC Genomics | 104 |
|  | Borowiec2015_Total1080 | Choaimalia | 1,080 | 384,981 | 36 | Ctenophora-sister |  |  |
| Pisani2015 | Ryan2013_est | Opisthokonta | 406 | 88,384 | 60 | Porifera-sister | PNAS | 206 |
|  | Whelan_D6 | Opisthokonta | 115 | 33,403 | 70 | Porifera-sister |  |  |
| Chang2015 | Chang2015 | Opisthokonta | 170 | 51,940 | 77 | Ctenophora-sister | PNAS | 119 |
| Whelan2016 | Philippe2009 | Opisthokonta | 129 | 30,257 | 55 | Ctenophora-sister | Systematic Biology | 37 |
|  | Nosenko2013_nonribo_9187 | Opisthokonta | 35 | 9,187 | 63 |  |  |  |
|  | Nosenko2013_ribo_14615 | Opisthokonta | 87 | 14,615 | 71 |  |  |  |
| Simion2017 | Simion2017 | Opisthokonta | 1,719 | 401,632 | 97 | Porifera-sister | Current Biology | 220 |
|  | Simion2017_subsampled | Opisthokonta | 1,719 | 110,602 | 97 | Porifera-sister |  |  |
| Feuda2017 | Chang2015 | Opisthokonta | 170 | 51,940 | 77 | Ctenophora-sister | Current Biology | 119 |
|  | WhelanD20 | Opisthokonta | 178 | 81,008 | 65 | Porifera-sister |  |  |
| Whelan2017 | Whelan2017b_full | Holozoa | 212 | 68,062 | 80 | Ctenophora-sister | Nature Ecology and Evolution | 79 |
|  | Whelan2017b_strict | Choaimalia | 117 | 49,388 | 76 | Ctenophora-sister |  |  |

## Extended Figures

**Extended Data Fig. 1.**
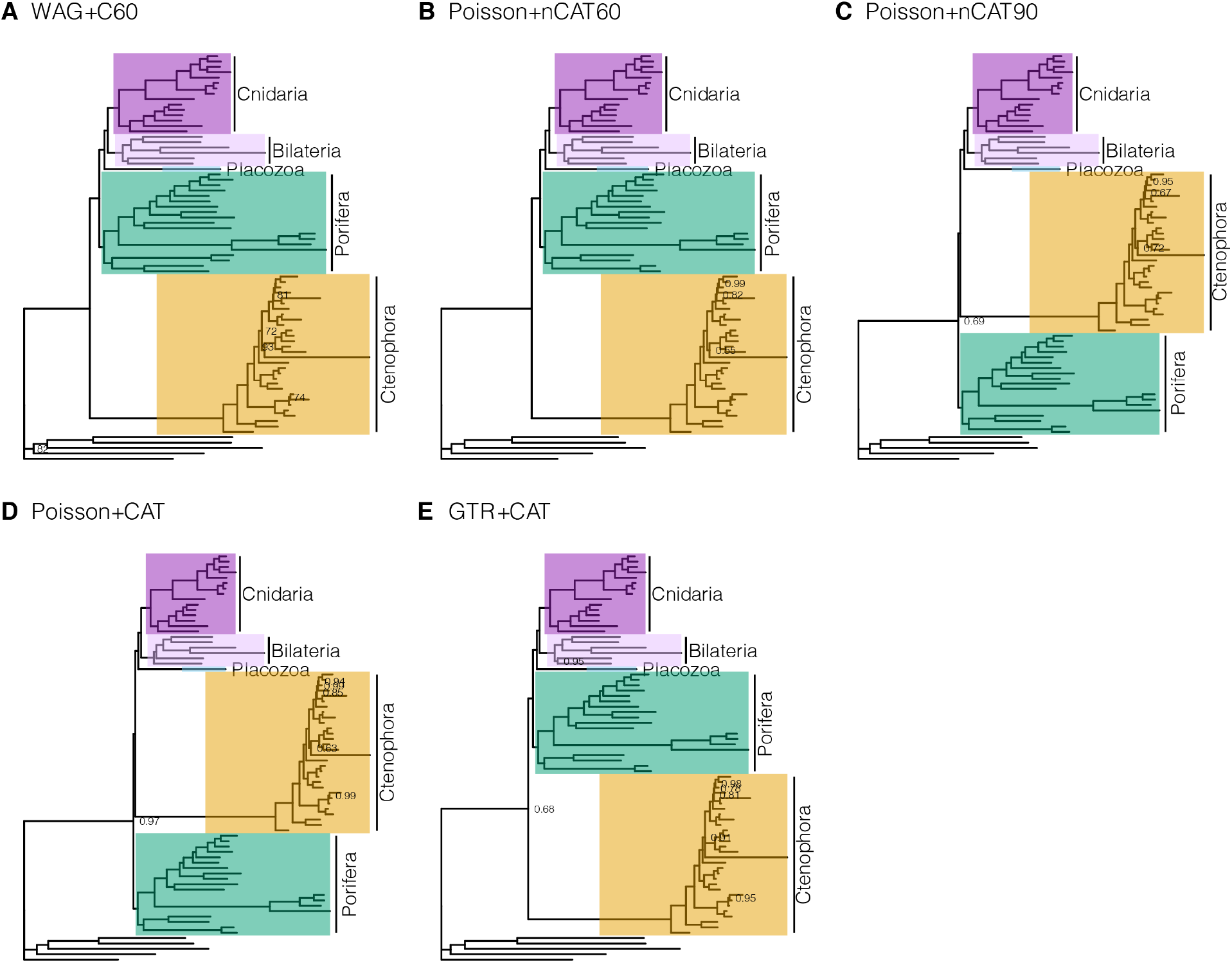
Comparison of the tree topologies under different substitutional models in Whelan2017_strict data matrix. (A). WAG+C60. (B). Poisson+nCAT60. (C). Poisson+nCAT90. (D). Poisson+CAT. (E). GTR+CAT. All trees were visualized and annotated in R package ggtree^30^.

**Extended Data Fig. 2.**
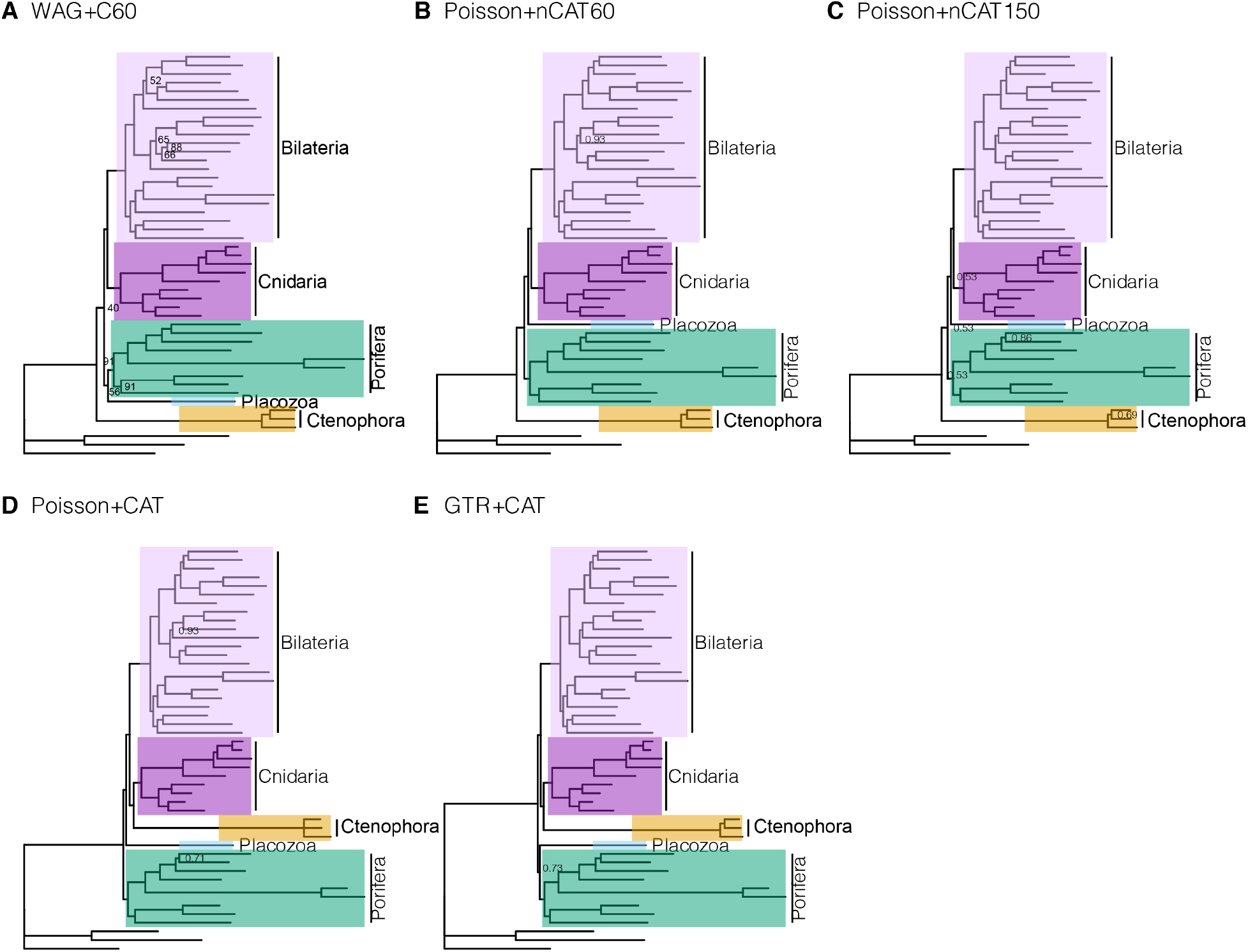
Comparison of the tree topologies under different substitutional models in Philippe2009_Choanozoa data matrix. (A). WAG+C60. (B). Poisson+nCAT60. (C). Poisson+nCAT150. (D). Poisson+CAT. (E). GTR+CAT. All trees were visualized and annotated in R package ggtree^30^.

**Extended Data Fig. 3.**
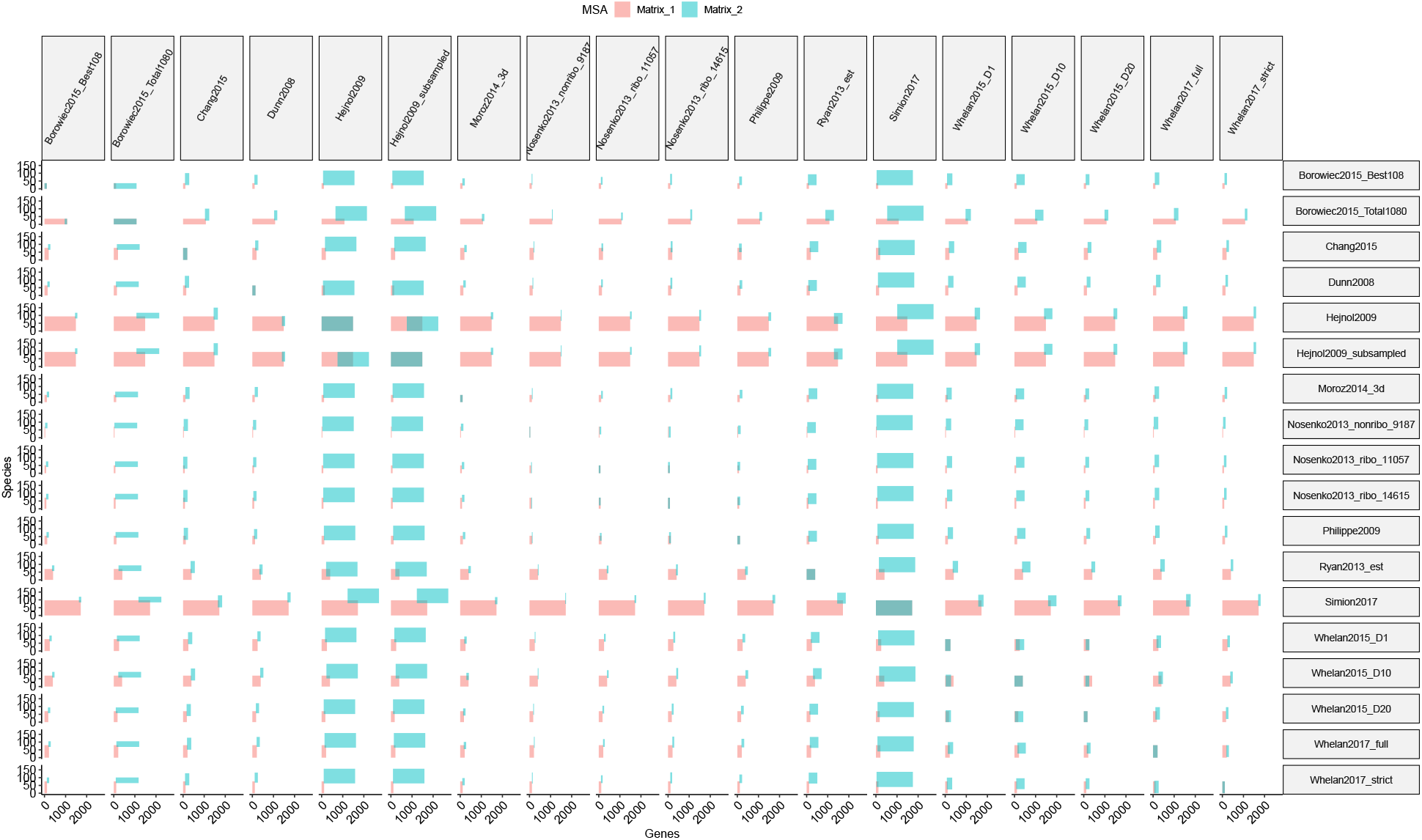
Pairwise overlap between each of the primary matrices considered here (related to Supplementary Table 8). Horizontal size is proportional to the number of genes sampled, vertical size to the number of taxa sampled. The horizontal intersection shows the proportions of shared genes, the vertical intersection shows the proportions of shared taxa.

**Extended Data Fig. 4.**
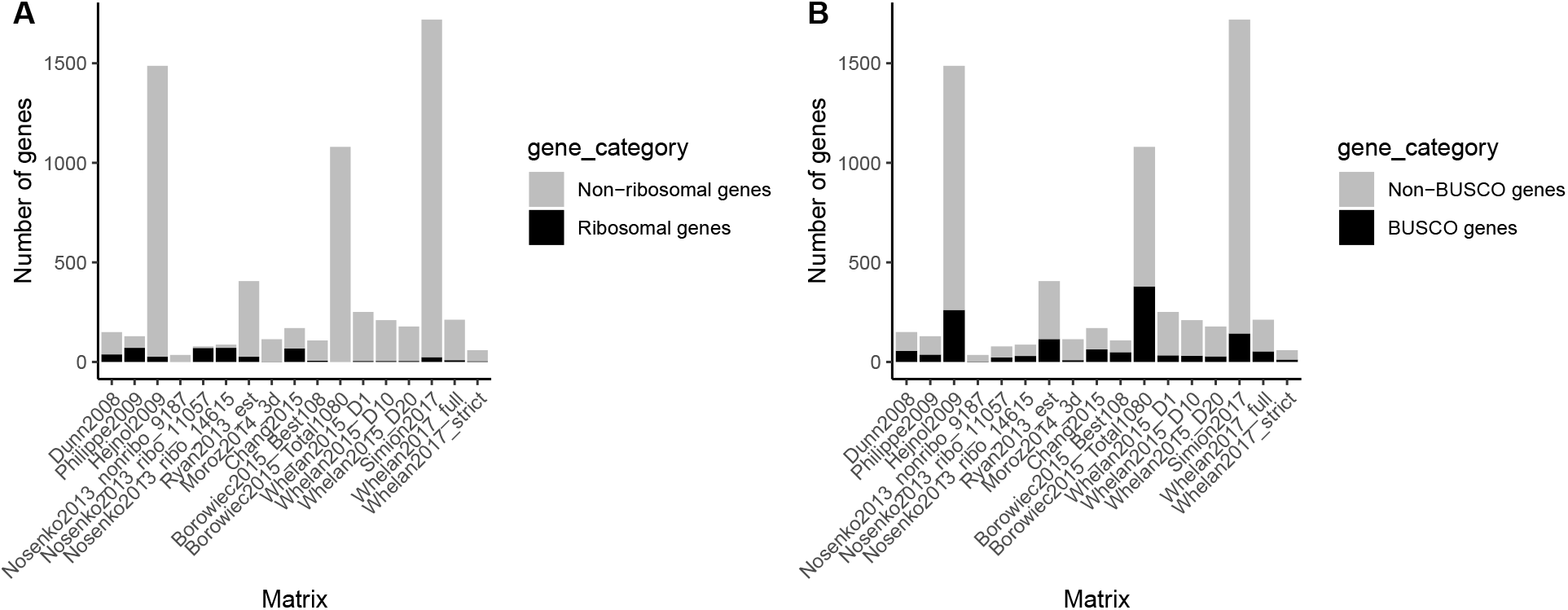
Annotation and representation of BUSCO and ribosomal genes in each data matrix (related to Supplementary Table 8). (A). The number of partitions with ribosomal annotations in each matrix, relative to the number of partitions. (C). The number of partitions with annotations of BUSCO genes in each matrix, relative to the number of partitions.

**Extended Data Fig. 5.**
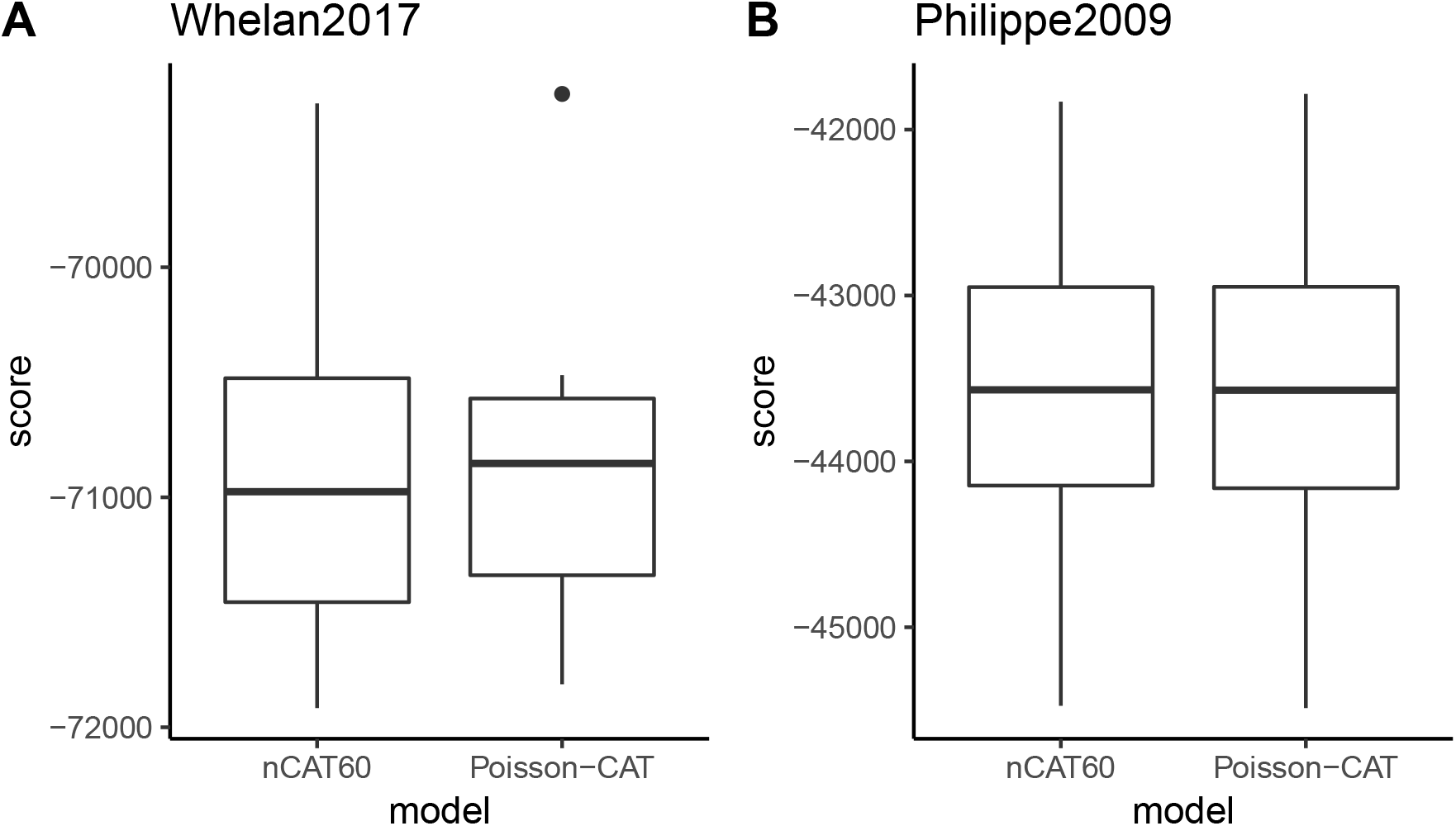
Box plots of log likelihood score for 10-fold cross-validation between Poisson+CAT and Poisson+nCAT60 models in Whelan2017_strict and Philippe2009_Choanozoa data matrices.

**Extended Data Fig. 6.**
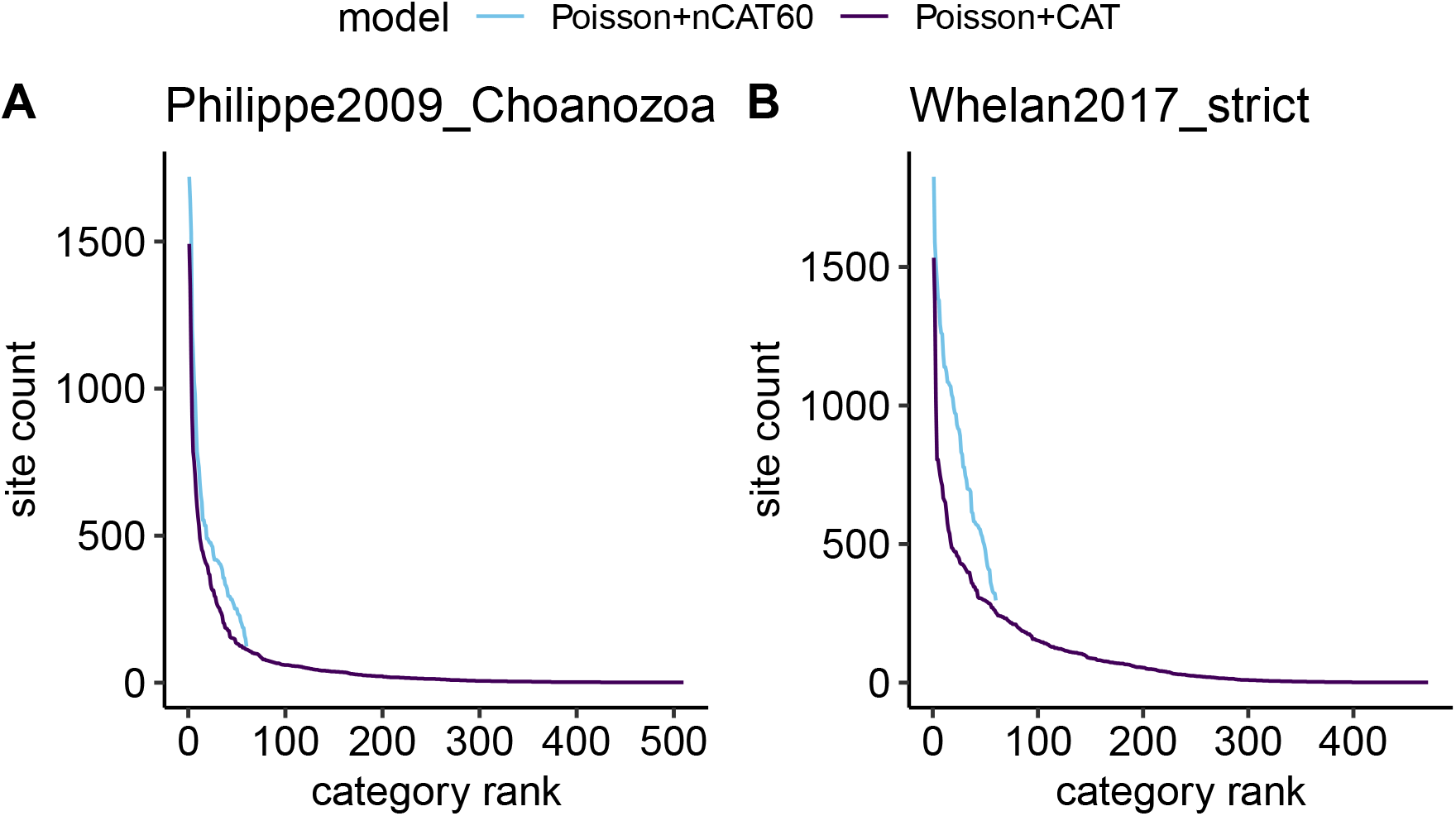
The allocation of frequency categories across sites in the Philippe2009_Choanozoa matrix (left column) and Whelan2017_strict matrix (right column) for the constrained Posson+nCAT60 model and unconstrained Posson+CAT models (differentiated with color). These plots are for singe sample isolated from the end of PhyloBayes chain files. (A,B) The count of sites allocated to each equilibrium frequency category, with the categories ranked from the most abundant to least abundant along the x axis. The unconstrained CAT analyses have a long tail of categories that are allocated to very few sites, which adds a considerable number of parameters that pertain to only a small fraction of the data. The nCAT60 analyses, which are constrained to 60 sites, have no such long tail and the rarest categories are allocated to a far greater number of sites than any of the sites in the long tail of the CAT analyses.

**Extended Data Fig. 7.**
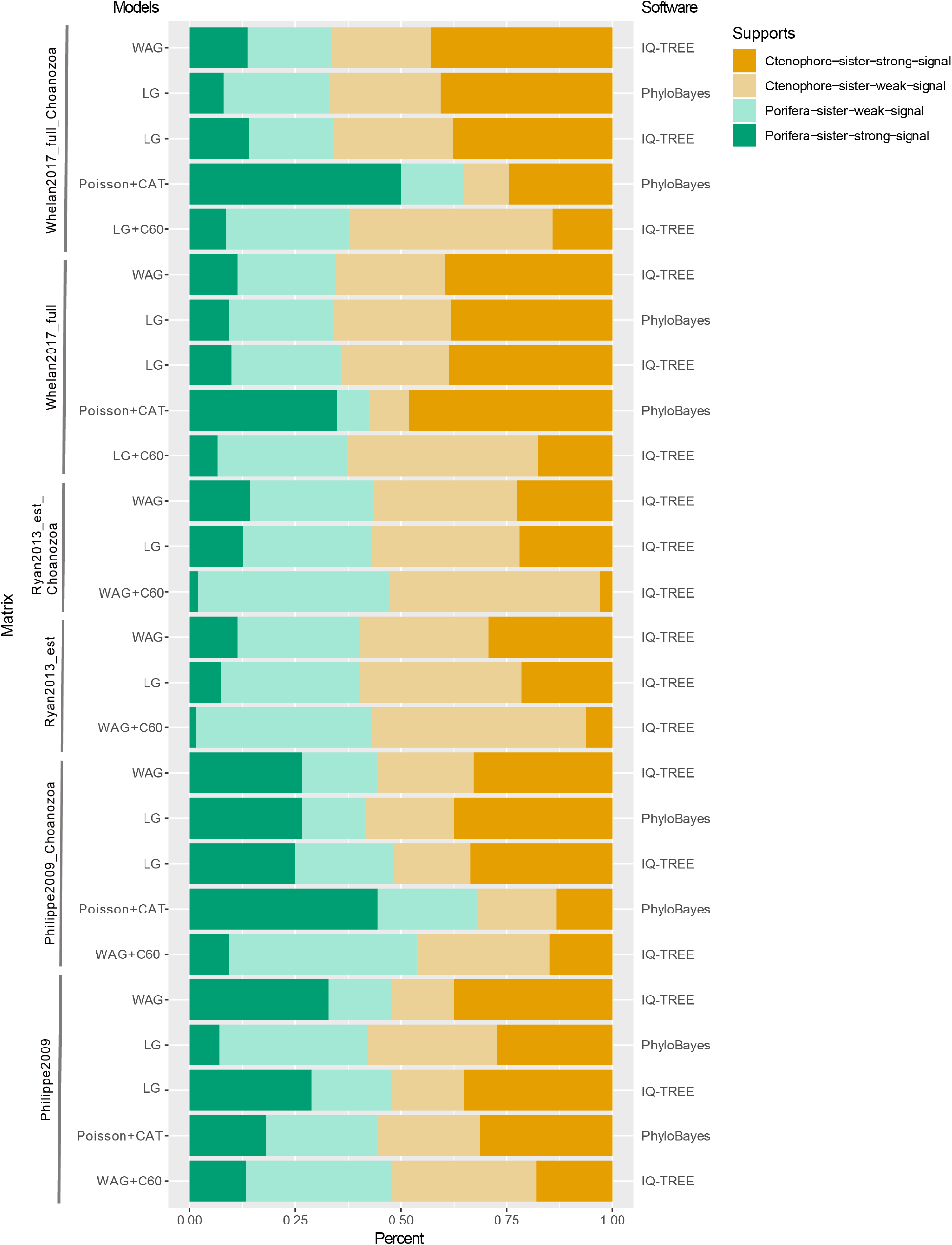
The distribution of phylogenetic signal for Ctenophora-sister and Porifera-sister with different models and outgroup choice in Philippe2009, Ryan2013 and Whelan2017_full matrices (with two outgroup sampling: Choanozoa and full matrices). The two alternative topological hypotheses are: Ctenophora-sister; T1 (Orange); Porifera-sister T2 (Green). Proportions of genes supporting each of two alternative hypotheses in the Philippe2009, Ryan2013_est and Whelan2017_full data matrices with different outgroups sampling and substitutional models. The GLS values for each gene in each data matrix are provided in Table S5. We considered a gene with an absolute value of log-likelihood difference of two as a gene with strong (|ΔlnL|< 2) or weak (0 < |ΔlnL|< 2) phylogenetic signal.

**Extended Data Fig. 8.**
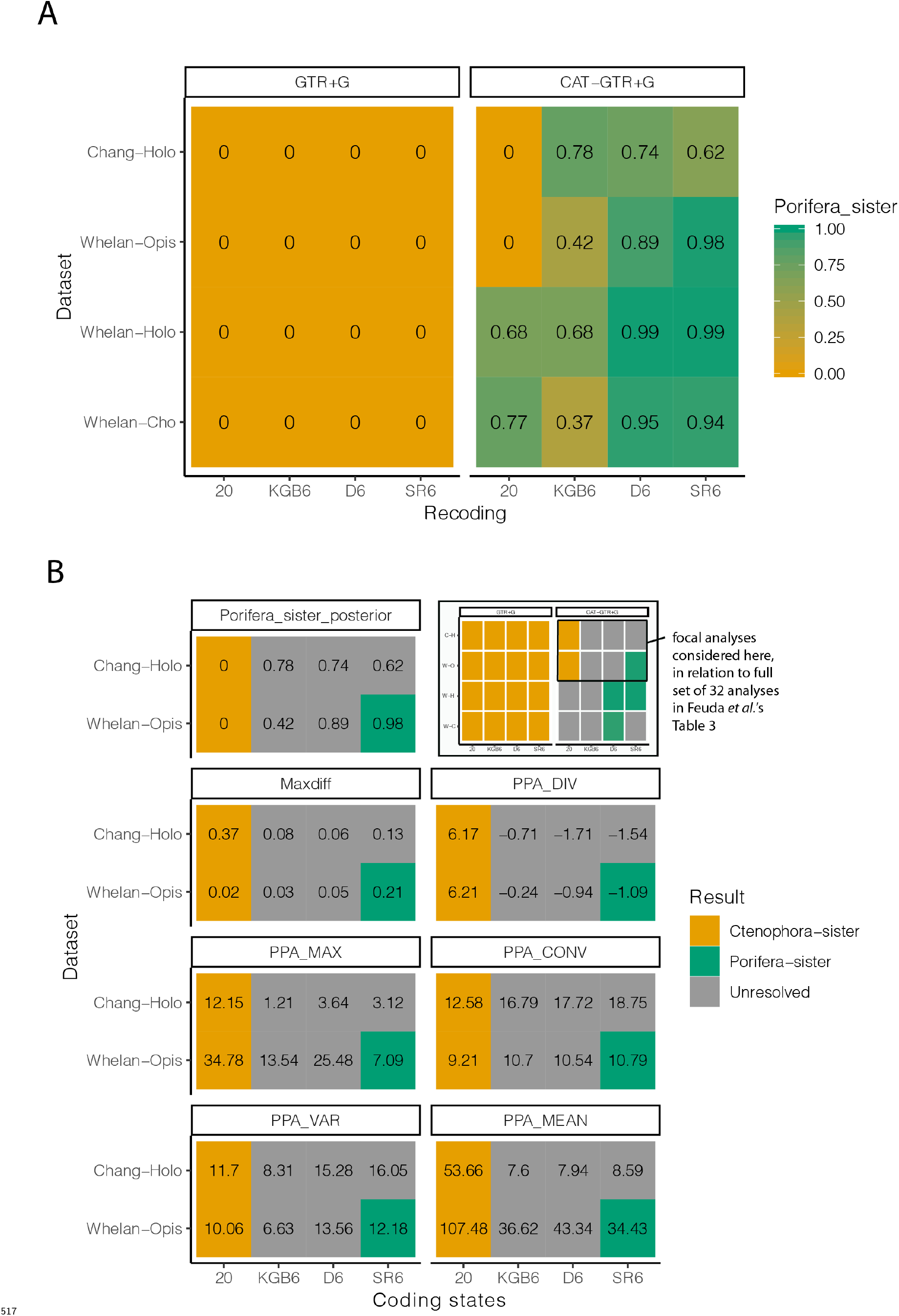
Graphical representations of the posterior probabilities for the 32 analyses presented by Feuda *et al.* in their Table 3 and the subset of eight GTR+CAT analyses with posterior predictive (PP) scores that is the focus of Feuda *et al.’s* primary conclusions. (A). Cells are color coded by whether posterior probability is > 95 for Porifera-sister, > 95 for Ctenophora-sister, or neither (unresolved). Posterior predictive (PP) statistics were estimated for the 16 analyses in the top two rows of this figure (the Chang and full Whelan matrices), but not the bottom two (the Whelan matrices with reduced outgroup sampling). (B). These are a subset of the 32 analyses presented in their Table 3 and graphically here in the upper right pane. The eight analyses are for two datasets (Chang and Whelan) and four coding schemes. The coding schemes are the original 20 state amino acid data, and three different six state recodings that group amino acids based on different criteria: KGB6, D6, and SR6. Only one of these analyses, the SR6 coding of the Whelan matrix, has > 95 support for Porifera-sister. The 20-state and 6-state points on the plots in Extended Data Fig. 8A correspond to the 20 and SR6 Whelan cells shown here. The presented statistics for these cells are posterior probability of Porifera-sister, Maxdiff (with lower scores indicating better convergence of runs), and five posterior predictive statistics (where lower absolute value indicates better model adequacy). The only one of these eight analyses that provides strong support for Porifera-sister is not the most adequate analysis by any of the posterior predictive scores, and showed the poorest convergence according to Maxdiff.

**Extended Data Fig. 9.**
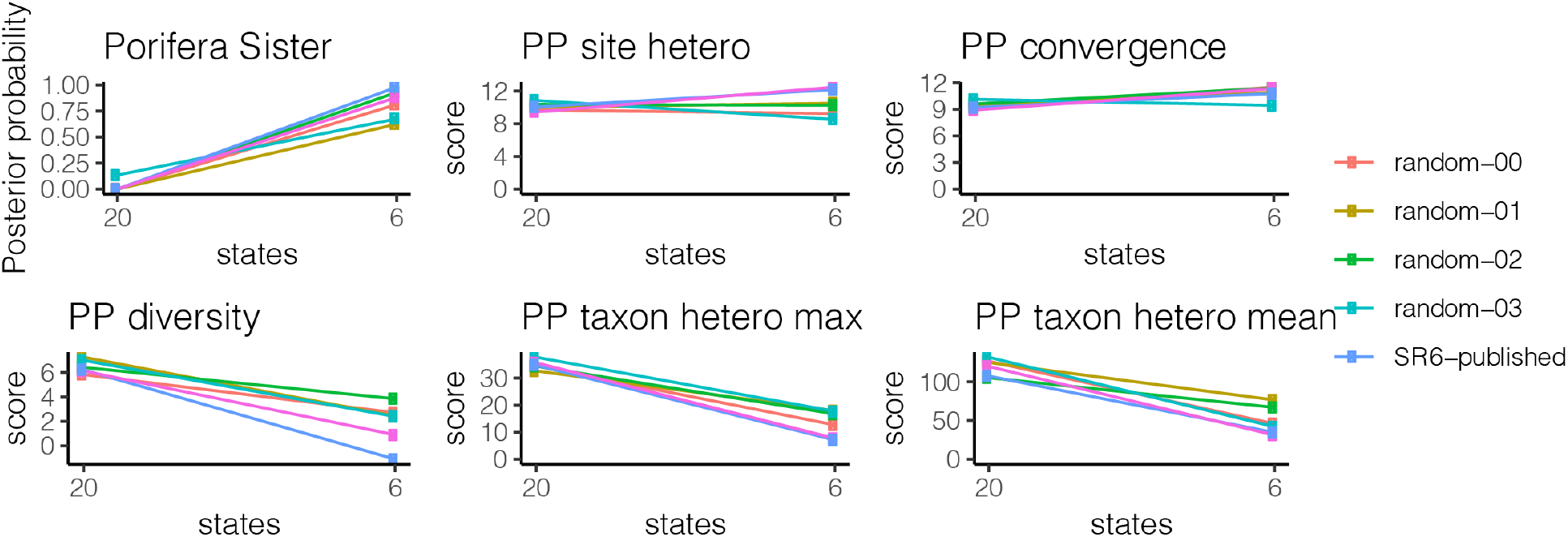
Each of the six plots presents one statistic, which include Posterior probability of Porifera-sister and the five Posterior Predictive (PP) statistics considered by Feuda *et al.* Within each plot, there are six lines for six different analyses. These five analyses are the published SR-6 analyses presented by Feuda *et al.* (SR6-published), and four analyses based on randomized recoding matrices obtained by shuffling the SR-6 coding scheme (random-00 – random-03). Each analysis includes results for 20 states (the raw amino acid data, shown by the left point) and for 6 states (the 6-state recoded data, shown by the right point). For each statistic, the results obtained with the random recoding are similar to those of the SR6 recoding. This indicates that the impact of recoding is dominated by discarding data when collapsing from 20 states to 6 states, not accommodating compositional hetezrogeneity across lineages.

## Supplementary Information

### S.1 Summaries of published analyses

Narrative summaries of the studies considered here.

#### Dunn *et al.* 2008

Dunn *et al.*^2^ added Expressed Sequence Tag (EST) data for 29 animals. It was the first phylogenomic analysis that included ctenophores, and therefore that could test the relationships of both Ctenophora and Porifera to the rest of animals. It was also the first phylogenetic analysis to recover Ctenophora as the sister group to all other animals.

The data matrix was constructed using a semi-automated approach. Genes were translated into proteins, promiscuous domains were masked, all gene sequences from all species were compared to each other with blastp, genes were clustered based on this similarity with TribeMCL^31^, and these clusters were filtered to remove those with poor taxon sampling and high rates of lineage-specific duplications. Gene trees were then constructed, and in clades of sequences all from the same species all but one sequence were removed (these groups are often due to assembly errors). The remaining gene trees with more than one sequence for any taxon were then manually inspected. If strongly supported deep nodes indicative of paralogy were found, the entire gene was discarded. If the duplications for a small number of taxa were unresolved, all genes from those taxa were excluded. Genes were then realigned and sites were filtered with Gblocks^32^, resulting in a 77 taxon matrix. Some taxa in this matrix were quite unstable, which obscured other strongly-supported relationships. Unstable taxa were identified with leaf stability indices^33^, as implemented in phyutility^34^, and removed from the matrix. This resulted in the 64-taxon matrix that is the focus of most of their analyses. Phylogenetic analyses were conducted under the Poisson+CAT model in PhyloBayes, and under the WAG model in MrBayes^35^ and RAxML^36^.

Regarding the recovery of Ctenophora-sister, the authors concluded:

> The placement of ctenophores (comb jellies) as the sister group to all other sampled metazoans is strongly supported in all our analyses. This result, which has not been postulated before, should be viewed as provisional until more data are considered from placozoans and additional sponges.

Note that there was, in fact, an exception to strong support. An analysis of the 40 ribosomal proteins in the matrix recovered Ctenophora-sister with only 69% support. This study did not include the placozoan ingroup *Trichoplax.*

#### Philippe *et al.* 2009

Philippe *et al.^16^* assembled an EST dataset for 55 species with 128 genes to explore the deepest animal phylogenetic relationships by adding 9 new species. The data matrix was assembled based on the phylogenetic analysis using Poisson+CAT model, which strongly supported Porifera-sister, and ctenophores were recovered as sister to cnidarians forming the “coelenterate” clade. Gene trees were then constructed, and potentially paralogs were removed by a bootstrap threshold of 70. Ambiguously aligned regions were trimmed and only genes sampled for at least two-thirds of species were retained. The phylogenetic analyses were conducted under the Poisson+CAT model in PhyloBayes.

Regarding the recovery of Ctenophora-sister, the authors concluded:

> The resulting phylogeny yields two significant conclusions reviving old views that have been challenged in the molecular era: (1) that the sponges (Porifera) are monophyletic and not pa– raphyletic as repeatedly proposed, thus undermining the idea that ancestral metazoans had a sponge-like body plan; (2) that the most likely position for the ctenophores is together with the cnidarians in a ‘coelenterate’ clade.

#### Hejnol *et al.* 2009

Hejnol *et al.^37^* added EST sequences from seven taxa, and a total of 94 taxa were included in the final data matrix to explore animal phylogeny, especially the position of acoelomorph flatworms. The orthology inference was largely similar to Dunn *et al.* 2008, with the exception of orthology genes which were clustered by MCL. The final data matrix included 1497 genes, and then subsampled with 844, 330 and 53 gens by different thresholds of gene occupancy. With the exception of 53 gene matrix, maximum likelihood analyses from all other datasets strongly supported Ctenophora-sister (models were selected by RaxML perl script).

#### Pick *et al.* 2010

Pick *et al.^3^* sought to test whether Ctenophora-sister was an artefact of insufficient taxon sampling. They added new and additional published sequence data to the 64-taxon matrix of Dunn *et al.^2^.* The new taxa included 12 sponges, 1 ctenophore, 5 cnidarians, and *Trichoplax.* They further modified the matrix by removing 2,150 sites that were poorly sampled or aligned. They considered two different sets of outgroups: Choanoflagellatea (resulting in Choanozoa) and the same sampling as Dunn *et al.* (resulting in Opisthokonta).

All their analyses were conducted under the F81+CAT+Gamma model in PhyloBayes, in both a Bayesian framework and with bootstrapping. All analyses have the same ingroup sampling and site removal so it isn’t possible to independently assess the impact of these factors. Analyses with Choanozoa sampling recovered Porifera-sister with 72% posterior probability (PP) and 91% bootstrap support (BS). With broader Opisthokonta sampling, support for Porifera-sister is 84% PP. This is an interesting case where increased outgroup sampling leads to increased support for Porifera-sister.

The authors argue that previous results supporting Ctenophora-sister “are artifacts stemming from insufficient taxon sampling and long-branch attraction (LBA)” and that “this hypothesis should be rejected”. Although the posterior probabilities supporting Porifera-sister are not strong, they conclude:

> Results of our analyses indicate that sponges are the sister group to the remaining Metazoa, and Placozoa are sister group to the Bilateria

They also investigated saturation, and concluded that the Dunn *et al.*^2^ data matrix is more saturated than Philippe *et al.* 2009 [Philippe:2009hh]. Note that the Pick *et al.^3^* dataset is not reanalyzed here because partition data are not available, and due to site filtering the partition file from Dunn *et al.*^2^ cannot be applied to this matrix.

#### Nosenko *et al.* 2013

Nosenko *et al^1^* added Expressed Sequence Tag (EST) data for 9 species of non-bilaterian metazoans (7 sponges). They constructed a novel matrix containing 122 genes and parsed them into two non-overlapping matrices (ribosomal and non-ribosomal genes) and found incongruent results of deep metazoan phylogeny. The other major finding was that ribosomal gene partitions showed significantly lower saturation than the non-ribosomal ones.

Orthologs were constructed using the bioinformatics pipeline OrthoSelect^38^. They also evaluated level of saturation, leaf stability of sampled taxa, compositional heterogeneity and model comparison of each matrix. By modifying gene sampling, ingroup and outgroup sampling, three major topologies related to the position of animal-root were constructed (including Porifera+Placozoa sister, Ctenophora-sister and Porifera-sister). Phylogenetic analyses were conducted under the Poisson+CAT, GTR+CAT and GTR models in PhyloBayes.

Regarding the recovery of Ctenophora-sister, the authors concluded:

> we were able to reconstruct a metazoan phylogeny that is consistent with traditional, morphology-based views on the phylogeny of non-bilaterian metazoans, including monophyletic Porifera and ctenophores as a sister-group of cnidarians.

#### Ryan *et al.* 2013

Ryan *et al.* sequenced the first ctenophore genome of *Mnemiopsis leidyi.* With the genome resources of *M. leidyi,* the authors constructed two phylogenomic datasets: a “Genome set” based on 13 animal genomes and a “EST Set” that also included 59 animals. They analyzed both matrices by site-homogeneous GTR+Gamma and site-heterogeneous Poisson+CAT models with three sets of outgroup sampling to evaluate the effect of outgroup selection to the ingroup topology for the Ryan2013_est matrix. The Orthologs were constructed based on the method of Hejnol *et al.* 2009. For the Ryan2013_genome matrix, they performed phylogenetic analyses with both gene content and sequence-based analyses. Overall, their results strongly supported Ctenophora-sister in all datasets they analyzed using a site-homogeneous model. The Poisson+CAT model of the genome dataset strongly supported of a clade of Ctenophora and Porifera as the sister group to all other Metazoa and Bayesian analysis on the EST dataset did not converge after 205 days (but strongly supported Porifera in Choaimalia matrix).

Regarding the recovery of Ctenophora-sister, the authors concluded:

> Our phylogenetic analyses suggest that ctenophores are the sister group to the rest of the extant animals.

#### Moroz *et al.* 2014

Moroz *et al.^39^* sequenced the second ctenophore genome *Pleurobrachia bachei* to explore the phylogenetic relationship of Metazoa. All phylogenetic analyses strongly supported Ctenophora-sister with different taxon and gene sampling using WAG site-homogeneous model. Two phylogenomic matrices were generated, the first set was represented by two ctenophore species, whereas the other set contained improved ctenophore sampling (10 taxa, Moroz2013_3d). Orthology determination employed in HaMStR^40^ using 1,032 “model organism” single-copy orthologs. Sequence were then trimmed and aligned. This resulted in a final matrix of 170,871 amino acid positions across 586 genes with 44 taxa for the first matrix, and 114 genes with 60 taxa for the second matrix. All the phylogenetic analyses were analyzed in RAxML under the WAG+CAT+F models (different from CAT models in PhyloBayes) to reduce the computational cost.

Regarding the recovery of Ctenophora-sister, the authors concluded:

> Our integrative analyses place Ctenophora as the earliest lineage within Metazoa. This hypothesis is supported by comparative analysis of multiple gene families, including the apparent absence of HOX genes, canonical microRNA machinery, and reduced immune complement in ctenophores.

It should be noted that only the Moroz_3d matrix has been reanalyzed in other studies, although the support of Ctenophora-sister is quite low.

#### Borowiec *et al.* 2015

Borowiec *et al.^41^* assembled a genome dataset comprising 1080 orthologs derived from 36 publicly available genomes representing major lineages of animals, although only one genome of sponge and ctenophore was included. The orthologs were constructed using OrthologID pipeline^42^. After removal of spurious sequences and genes with more than 40% of mission data, the final matrix included 1080 (Total 1080) genes. The authors further filtered the full dataset to 9 sub-datasets by filtering genes with high long-branch scores; genes with high saturation; gene occupancy; fast evolving genes. The main conclusion of the study was largely based on BorowiecTotal_1080 and Borowiec_Best108 matrices. Phylogenetic analyses were conducted under the GTR+CAT model in PhyloBayes in selected matrices, and under the data-partitioning methods in RAxML for all matrices.

Regarding the recovery of Ctenophora-sister, the authors concluded:

> Our phylogeny supports the still-controversial position of ctenophores as sister group to all other metazoans. This study also provides a workflow and computational tools for minimizing systematic bias in genome-based phylogenetic analyses.

It should be noted that the authors also employed recoding-method in the Borowiec_Best108 matrix and found neither support of Porifera-sister or Ctenophora-sister^41^.

#### Whelan et al. 2015

Whelan *et al.* 2015^43^ constructed a new phylogenomic data matrix with eight new transcriptomic data and investigated a range of possible sources of systematic error under multiple analyses *(e.g.* long-branch attraction, compositional bias, fast evolving genes, etc.). Putative orthologs were determined of each species using HaMStR using the model organism core ortholog set (same as Moroz *et al.* 2014) and subsequently removal of genes with too much missing data and potential paralogs. The authors further filtered the full dataset to 24 sub-datasets by filtering genes with high long-branch scores; genes with high RSFV values; genes that are potential paralogs; fast evolving genes and progressive removal of outgroups. All the maximum likelihood analyses with site-homogeneous models and PartitionFinder strongly suggested Ctenophora-sister. GTR+CAT models only used in slow-evolving data matrices 6 and 16 also strongly supported Ctenophora.

Regarding the recovery of Ctenophora-sister, the authors concluded:

> Importantly, biases resulting from elevated compositional heterogeneity or elevated substitution rates are ruled out. Placement of ctenophores as sister to all other animals, and sponge monophyly, are strongly supported under multiple analyses, herein.

Note that the authors also reanalyzed Philippe2009 matrix (with the removal of ribosomal genes) and recovered Porifera-sister with moderate support (pp=90).

#### Chang et al. 2015

Chang *et al.^44^* was originally used to explore the phylogenetic position of Myxozoa in Cnidaria but also sampled broadly across the breadth of animal diversity. The authors constructed a dataset with 200 protein markers based on Philippe *et al.* 2011^45^ with 51,940 amino acids and 77 taxa. Both site-heterogeneous Poisson+CAT and site-homogeneous GTR models strongly supported Ctenophora-sister.

#### Pisani *et al.* 2015

Pisani *et al.^5^* reanalyzed representative datasets that supported Ctenophora-sister, including Ryan2013_est, Moroz2014_3d and Whelan2015 datasets. It was the first study showing that progressive removal of more distantly related outgroups could largely affect phylogenomic inference of the position of the root of animal phylogeny. The authors suggested that the inclusion of outgroups very distant from the ingroup can cause systematic errors due to long-branch attraction. Phylogenetic analyses were conducted under the Poisson+CAT and GTR models in PhyloBayes. They found Poisson+CAT models generally had better model-fit than site-homogeneous GTR models in these data matrices. Moreover, they found the support of Ctenophora-sister decreases when the exclusion of distantly related outgroups are excluded and the use of site-heterogeneous CAT models are used.

Regarding the recovery of Porifera-sister, the authors concluded:

> Our results reinforce a traditional scenario for the evolution of complexity in animals, and indicate that inferences about the evolution of Metazoa based on the Ctenophora-sister hypothesis are not supported by the currently available data.

#### Feuda *et al.* 2017

Feuda *et al^7^* didn’t generate any new data, instead they used the data-recoding methods to reanalyze two key datasets that support Ctenophora-sister (Whelan2015_D20, Chang2015 datasets). It was the first phylogenomic study that suggested recoding methods have better performance than non-recoding methods based on recovering Porifera-sister hypothesis. The authors compared model adequacy using posterior predictive analyses from a set of site-homogeneous (WAG, LG, GTR, data-partitioning) and site-heterogeneous (GTR+CAT) models in non-recoding and recoding datasets. The results showed that data-recoding can significantly reduce compositional heterogeneity in both datasets with GTR+CAT models and strongly supported Porifera-sister hypothesis (see more details in Supplementary Information section S3).

Regarding the recovery of Porifera-sister, the authors concluded:

> Because adequate modeling of the evolutionary process that generated the data is fundamental to recovering an accurate phylogeny, our results strongly support sponges as the sister group of all other animals and provide further evidence that Ctenophora-sister represents a tree reconstruction artifact.

#### Whelan and Halanych 2016

Whelan *et al.^6^* is the only study to evaluate performance of site-heterogeneous models and site-homogeneous models with data partitioning under the simulation framework. The simulation results suggested that the Poisson+CAT model consistently performed worse than other models in simulation datasets. More importantly, the authors also showed that both Poisson+CAT and GTR+ CAT models could overestimate substitutional heterogeneity in almost every case. They also reanalyzed datasets from Philippe2009 and Nosenko2013 using both CAT models and data partitioning with site-homogeneous model. The results indicated that Poisson + CAT model tends to recover less accurate trees and both GTR + CAT and data partitioning strongly supported Ctenophora-sister in reanalyses.

The authors concluded:

> Practices such as removing constant sites and parsimony uninformative characters, or using CAT-F81 when CAT-GTR is deemed too computationally expensive, cannot be logically justified. Given clear problems with CAT-F81, phylogenies previously inferred with this model should be reassessed.

#### Whelan *et al.* 2017

Whelan *et al.^46^* added 27 new ctenophore transcriptomic data to explore animal-root position as well as relationships within Ctenophores. It significantly increased ctenophore taxon sampling than other studies. Putative orthologs were determined largely similar to Whelan2015. The subsequent filtering strategy was also similar to the previous study. All analyses using site-homogeneous and site-heterogeneous models strongly supported Ctenophora-sister hypothesis, even with GTR+CAT model in Choanozoa dataset. The main conclusions of this study were based on Whelan2017_full and Whelan2017_strict matrices.

Regarding the recovery of Ctenophora-sister, the authors concluded: >Using datasets with reasonably high ctenophore and other non-bilaterian taxon sampling, our results strongly reject the hypothesis that sponges are the sister lineage to all other extant metazoans.

#### Simion *et al.* 2017

Simion *et al*.^18^ added transcriptomic data for 21 new animals. The data matrix was constructed using a semi-automated approach to comprehensively detect and eliminate potential systematic errors. The resulting dataset comprises 1,719 genes and 97 species, including 61 non-bilaterian species. It was by far the largest phylogenomic dataset in terms of taxon and gene sampling related to the relationship at the root of animal phylogeny.

The final matrix was first analyzed using the Poisson+CAT model. Different from other PhyloBayes analyses, Simion *et al.* used a gene jackknife strategy based on 100 analyses to overcome the computational limitation because of the large data size. Each jackknife is based on a random selection of ~ 25% of the genes. The PhyloBayes with site-heterogeneous model strongly supported the Porifera-sister, whereas site-homogeneous strongly supported Ctenophora-sister in all datasets. Importantly, the authors compared the behavior of long-branch effect between site-heterogeneous and site-homogeneous models by progressively removing taxa and concluded higher sensitivity of site-homogeneous models to LBA than CAT models.

Regarding the recovery of Ctenophora-sister, the authors concluded:

> Our dataset outperforms previous metazoan gene super alignments in terms of data quality and quantity. Analyses with a best-fitting site-heterogeneous evolutionary model provide strong statistical support for placing sponges as the sister-group to all other metazoans, with ctenophores emerging as the second-earliest branching animal lineage.

It should be noted that all the PhyloBayes runs have not reached convergence due to the computational cost in these large matrices.

### S.2 Models of molecular evolution

The exchangeability matrix *R* describes the relative rates at which one amino acid changes to others. Exchangeability matrices have been used in the studies under consideration here include: Poisson (or F81), WAG, LG, GTR. While the exchangeability matrix describes the relative rate of different changes between amino acids, the actual rate can be further scaled. There are a couple approaches that have been used in the studies considered here:

- F81^47^ corresponds to equal rates between all states. The F81 matrix is also sometimes referred to as the Poisson matrix. It has no free parameters to estimate since all off-diagonal elements are set to 1.
- WAG^48^ is an empirically derived exchangeability matrix based on a dataset of 182 globular protein families. It has no free parameters to estimate since all off-diagonal elements are set according to values estimated from this particular sample dataset.
- LG^49^, like WAG, is an empirically derived exchangeability matrix. It is based on a much larger set of genes, and variation in rates across sites was taken into consideration when it was calculated. It has no free parameters to estimate since all off-diagonal elements are set according to values estimated from this particular sample dataset.
- GTR, the General Time Reversible exchangeability matrix, has free parameters for all off-diagonal elements that describe the exchangeability of different amino acids. It is constrained so that changes are reversible, *i.e.* the rates above the diagonal are the same as those below the diagonal. This leaves 190 parameters that must be estimated from the data along with the other model parameters and the phylogenetic tree topology. This estimation requires a considerable amount of data and computational power, but if successful has the advantage of being based on the dataset at hand rather than a different dataset (as for LG and WAG).

While the exchangeability matrix describes the relative rate of different changes between amino acids, the actual rate can be further scaled. There are a couple approaches that have been used in the studies considered here:

- Site homogeneous rates. The rates of evolution are assumed to be the same at all sites in the amino acid alignment.
- Gamma rate heterogeneity. Each site is assigned to a different rate class with its own rate value. This accommodates different rates of evolution across different sites. Gamma is used so commonly that sometimes it isn’t even specified, making it difficult at times to know if a study uses Gamma or not.

The vector of equilibrium frequencies Π describes the stationary frequency of amino acids. There are a few approaches that have been used across the studies considered here:

- Empirical site-homogeneous. The frequency of each amino acid is observed from the matrix under consideration and applied to homogeneously to all sites in the matrix.
- Estimated site-homogeneous. The frequency of each amino acid is inferred along with other model parameters, under the assumption that it is the same at all sites.
- CAT site-heterogeneous. Each site is assigned to a class with its own equilibrium frequencies. The number of classes, assignment of sites to classes, and equilibrium frequencies within the data are all estimated in a Bayesian framework.
- C10 to C60^14^. 10 to 60-profile mixture models as variants of the CAT model under the maximumlikelihood framework.

### S.3 Data-recoding methods

Feuda *et al.^7^* were concerned that the Ctenophora-sister results Chang *et al.^44^* and Whelan *et al.^50^* were artefacts of lineage-specific differences in amino acid frequencies. In an attempt to reduce these differences, they recoded the full set of twenty amino acids into six groups of amino acids. These groups have more frequent evolutionary changes within them than between them, based on empirical observations in large protein datasets^22^. The intent is to discard many lineage-specific changes, which are expected to fall within these groups. Rather than model compositional heterogeneity, as their title suggests, this approach discards heterogeneous information so that much simpler models with fewer states can be applied.

Feuda *et al.^7^* report that posterior predictive (PP) analyses^51^ indicate 6-state recoded analyses have better model adequacy than 20-state amino acid analyses, and “Porifera-sister was favored under all recoding strategies” in Whelan2015_D20 and Chang2015 data matrices. Here we focus on two aspects of Feuda *et al.* First, we point out that many of their recoded analyses are actually unresolved *(i.e.,* without strong support for either Porifera-sister or Ctenophora-sister) (Extended Data Fig. 8A). Second, we present new analyses that show the impact of recoding is largely due to discarding information, not accommodating variation in amino acid composition. These findings indicate that recoding can be a problematic method for addressing compositional variation.

Feuda *et al.* examine support for Ctenophora-sister and Porifera-sister under all combinations of two models of molecular evolution, four datasets, and four coding schemes. This provides 32 analyses that they report in their Table 3 and that we present graphically here as Extended Data Fig. 8A. There is striking variation in support for Ctenophora-sister and Porifera-sister across these analyses (Extended Data Fig. 8A). Feuda *et al.* accept the results of some analyses and reject others based on posterior predictive (PP) analyses of model adequacy, which assess how well a model explains variation in the data^51^. They considered five different posterior predictive statistics that capture different types of variation in the data. From this they conclude that their “results strongly support sponges as the sister group of all other animals”.

This conclusion does not follow from their own presented results. Only a single analysis with posterior predictive scores provides what could be considered strong support > 95 posterior probability) for Porifera-sister. Of the 32 analyses, posterior predictive scores were calculated for 16 (those for the full Whelan and Chang matrices). Based on posterior predictive scores, Feuda *et al.* reject eight of these that were conducted under the GTR+G model (which all have strong support for Ctenophora-sister). This leaves eight GTR+CAT analyses (Extended Data Fig. 8A). Two of these eight are analyses of the original 20-state amino acid data, both of which provide strong support for Ctenophora-sister. Of the six recoded analyses, five are unresolved. Only a single analysis for which posterior predictive scores are available provides strong support for Porifera-sister, the GTR+CAT analysis of the SR-6^22^ recoded Whelan^50^ matrix. Furthermore, this analysis does not have the best score according to any of the five posterior predictive statistics they considered (Extended Data Fig. 8B). The only statistic that stands out for this one analysis is that it has the highest maxdiff (Extended Data Fig. 8B), indicating that it did not converge as well as other analyses.

Though their study does not provide strong support for Porifera-sister, the sensitivity of their results to recoding provides an opportunity to better understand and evaluate the impact of recoding more generally. This is important given the growing interest in recoding^21^. Feuda *et al.* hoped recoding would reduce potential artefacts due to differences across species in amino acid frequencies. They interpreted the fact that their analyses are sensitive to recoding as evidence that such an artefact exists and that they successfully addressed it by recoding. An alternative hypothesis is that recoding impacts phylogenetic analyses because it discards so much information. These two hypotheses can be tested by applying new recoding schemes that also reduce twenty states down to six, but are based on random grouping rather than empirical frequencies of amino acid exchange. Empirical and random recodings both discard the same amount of information, but only empirical recoding reduce the impact of amino-acid frequency as intended. Different results between empirical and random recoding would be consistent with the hypothesis that the empirical approach works as intended to accommodate compositional heterogeneity. Similar results would suggest that the impact of recoding is due to discarding information. Here we focus on their single analysis with a posterior predictive score that supports Porifera-sister, the GTR+CAT analysis of the SR-6 recoded Whelan data. We created four new random recoding schemes by shuffling the amino acids in the SR-6 scheme (see Supplemental Methods and analysis code at https://github.com/caseywdunn/feuda_2017). When we applied each of these randomized codes to the Whelan matrix and analyzed them under the GTR+CAT model with PhyloBayes-MPI, we observed similar results as for the empirical SR-6 recoding. Like SR-6 recoding, random recoding increases support for Porifera-sister and improves the apparent adequacy of models to explain heterogeneity of states across taxa (PP taxon hetero mean and max, Extended Data Fig. 9).

These analyses suggest that the major impact of recoding on phylogenetic analyses is data reduction, not accommodation of compositional heterogeneity across species. This indicates that recoding may not be an effective tool for accommodating among-species differences in amino acid frequencies. Compositional heterogeneity would be better addressed with models of molecular evolution that explicitly describe such differences^52^, if progress can be made on the considerable computational challenges of such complex models.

### Supplementary Tables

**Supplementary Table 1.** Summary of a total of 164 phylogenomic analyses were transcribed from the literature (Table is converted from analyses_published in Rdata).

**Supplementary Table 2.** Summary of a total of 106 phylogenomic analyses conducted in this study (Table is converted from analyses_new in R data).

**Supplementary Table 3.** The models selected by ModelFinder for each matrix in IQ-TREE.(Table is converted from analyses_new in Rdata).

**Supplementary Table 4.** Summary statistics of CAT substitutional categories inferred from different matrices (Table is manualy curated from Tracer result for each PhyloBayes analysis).

**Supplementary Table 5.** Summary of sensitive analyses with different number of CAT substitutional categories in representative matrices (Table is converted from analyses_sensitive in Rdata).

**Supplementary Table 6.** Summary of amino acid frequencies of 60 categories inferred by C60 model using IQ-TREE (Table is manualy curated from IQtree log file for each analysis).

**Supplementary Table 7.** Distribution of phylogenetic signal of different models and outgroup sampling for two alternative hypotheses on the animal-root position in Philippe2009, Ryan2013 and Whelan2017_full matrices (Table is converted from au_tests in Rdata).

**Supplementary Table 8.** Summary of annotations (BUSCO, Ensemble, GO-terms, ribosomal protein genes) and network analyses for each partition from all phylogenomic all matrices used in this study (Table is converted from partition_map_globle in Rdata).

